# Bidirectional Control of Parathyroid Hormone and Bone Mass by Subfornical Organ

**DOI:** 10.1101/2020.11.30.403279

**Authors:** Lu Zhang, Nian Liu, Jie Shao, Dashuang Gao, Yunhui Liu, Di Chen, Liping Wang, William Weijia Lu, Fan Yang

## Abstract

Parathyroid hormone (PTH) is one of the most important hormones responsible for bone turnover and calcium homeostasis. In mammals, PTH is secreted through the parathyroid glands, unlike in fishes, which are secreted through central nervous system. Studies indicated that a variety of peripheral nerve regulates serum PTH level, however, the mechanism underlying central neural regulation of PTH in mammals remains largely unknown. With an approach including neural-specific retrograde tracing, PTH-Biotin binding assay, and cFos staining, we identified the subfornical organ (SFO) and the paraventricular nucleus (PVN) as two important brain nuclei that responded to serum PTH and calcium changes. Using chemogenetics, we found that serum PTH was suppressed by stimulation of GABAergic neurons in SFO followed by a decrease in trabecular bone mass. Conversely, stimulation of glutamatergic neurons in SFO promoted serum PTH and bone mass. Moreover, we found that the majority of neurons expressing parathyroid hormone 1 receptor (PTH1R) were GABAergic neurons and the majority of neurons expressing parathyroid hormone 2 receptor (PTH2R) were glutamatergic neurons. The paraventricular nucleus (PVN) is downstream of the SFO, and chemogenetic activation of glutamatergic neurons in PVN induced an increase in serum PTH. In summary, our study demonstrates for the first time that distinct neuronal subtypes in the SFO are responsible for bidirectional regulation of serum PTH and bone metabolism, which is mediated through the PVN and the peripheral nervous system. These findings reveal important central neural nodes and will advance our understanding of the central neural regulation of PTH at the molecular, cellular and circuit level.

## Introduction

Parathyroid hormone (PTH) is one of the most important hormones that modulate bone remodeling and calcium homeostasis, which is conserved from zebrafish to humans^1,2^. In fish that do not have isolated parathyroid glands, PTH peptides are mainly expressed in the lateral line, the neural tube and in the central nervous system^3–5^. In humans and other mammals, PTH is mainly secreted from the parathyroid glands to maintain basal serum calcium levels under different nutrient conditions. The secretion of PTH is finely regulated by the humoral, hormonal stimulations, and the level of PTH is precisely maintained based on the serum calcium concentration.

In the mammalian central nervous system (CNS), parathyroid hormone 2 receptors (PTH2R) and TIP39 proteins form a unique neuropeptide-receptor system, in which there is a wide distribution of axon terminals from neurons that support functions such as nociceptive signaling and neuroendocrine regulation^6^. In the rat brain, PTH2-receptor expression is observed in the cerebral cortex, stratum and hypothalamus, including the arcuate nuclei and median eminence^7^. Recent work demonstrated that ancient parathyroid hormone 4 (PTH4) appear to be a central neural regulator of bone development and mineral homeostasis^5^, and ablation of PTH4 neurons results in abnormal bone mineralization and osteoblast differentiation in zebrafish^8^. These findings support that PTH or PTH receptors might have played a continuous prominent role in the brain-to-bone signaling pathway and in the maintenance of bone homeostasis during vertebrate evolution. However, whether the adult mammalian central nervous system has retained the capacity to detect PTH levels and direct the neural regulation of PTH homeostasis is largely unknown.

In addition to the CNS, abundant levels of neural terminal expression have been observed in parathyroid glands^9–14^. Administration of epinephrine or isoproterenol infusion acutely induces PTH release, which is ablated by the β-adrenergic receptor 2 antagonist propranolol^15^, and direct stimulation of parathyroid nerves upstream of the parathyroid induces serum PTH elevation^16^. Clinical studies have also shown that neuropsychiatric symptoms including anxiety, depression and cognitive impairments are prevalent in patients with hyperparathyroidism, which, along with bone abnormalities, are significantly improved after parathyroidectomy^17,18^. Based on this evidence, we hypothesized that distinct brain nuclei and neuronal subtypes may sense PTH levels and direct peripheral nerves to regulate PTH secretion in a top-down manner. However, it is not clear what these important nodes in the central nervous system that control PTH secretion in mammals are, or the underlying mechanisms.

In this study, we firstly identify the subfornical organ (SFO), a member of the circumventricular organs (CVO), as an important brain nucleus modulating the homeostasis of PTH. Chemogenetic activation of glutamatergic and GABAergic neurons in SFO exerted responses in opposing directions in serum PTH and tibiae trabecular bone mass. Neurons in the SFO express PTH receptors 1 and 2, the majority of which are expressed in GABAergic and glutamatergic neurons, respectively. Activation of glutamatergic nerves in paraventricular nucleus (PVN), which are downstream of the SFO, led to elevated serum PTH and trabecular bone mass. In addition, sympathetic and sensory nerves were identified within the parathyroid glands, and ablation of these affected serum PTH levels and PTH response to calcium. Our study not only demonstrates that the central nervous system is indispensable for the regulation of peripheral PTH secretion and maintenance of bone-metabolism homeostasis, but also for the first time revealed the underlying mechanism of central neural regulation of the PTH at molecular, cellular and circuit level.

## Results

### Parathyroid glands are connected to the central nervous system

Previous studies have demonstrated that parathyroid glands (PTG) receive sympathetic, parasympathetic, and sensory neural innervation, however, the central source of this innervation was not clear. To determine the brain nucleus that innervates the parathyroid gland (PTG), we injected the neural retrograde tracer pseudorabies virus (PRV-GFP) directly into the PTG (Fig. 1A-F). It could be observed the pseudorabies virus infects the central nervous system through the brain stem initiating from the intermediate reticular nucleus (IRt) and raphe magnus nucleus (RMg) (Fig. 1C-D). From the date of virus administration, it usually takes 7 days for the virus to infect from brainstem to midbrain and then to the hypothalamus and forebrain (Fig. 1E). The brain nuclei with the highest PRV expression markers on the 7^th^ day of infection included: the median preoptic nucleus (MnPO), subfornical organ (SFO), suprachiasmatic nucleus (SCh), the nucleus of the vertical limb of the diagonal band (VDB), and Arcuate nucleus (Arc) (Fig. 1F, Supplementary Fig. 1). The MnPO and SFO and area postrema (AP) belongs to a group called the circumventricular organs (CVO), which are characterized by their extensive and highly permeable microvasculature and have no blood-brain-barrier^19^.

**Figure 1.**
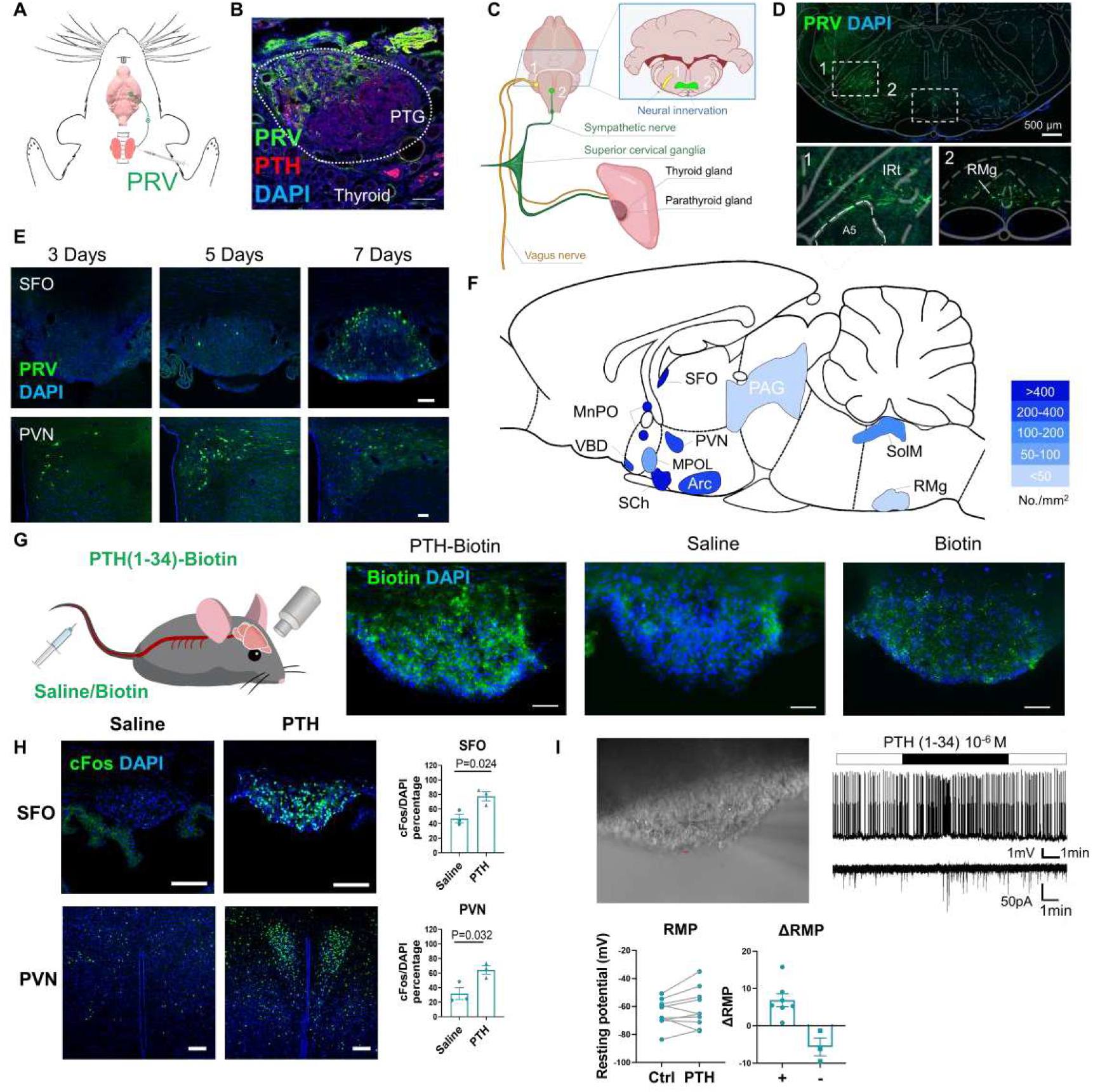
Parathyroid glands are connected to the SFO and PVN. **A**, Schematic showing pseudorabies virus (PRV) infection of SD rat parathyroid gland (PTG) and poly trans-synaptic connection to the central nervous system. **B,** PRV *in situ* expression within rat PTG. Scale bar, 50 μm. **C**, Schematic showing the peripheral neural innervation of the parathyroid glands and the connection ends to brain stem. **D,** PRV expression in brain stem 3 days after administration to the rat parathyroid gland. IRt, intermediate reticular nucleus; RMg, raphe magnus nucleus. **E**, PRV expression in subfornical organ (SFO, ***upper,*** scale bar, 100 μm) and paraventricular nucleus (PVN, ***lower,*** scale bar, 100 μm) at 3,5 and 7 days after injection into the rat parathyroid gland. **F**, PRV expression density in different rat brain areas after PRV neural tracing (No./mm^2^ n=3). Arc, arcuate nucleus; MnPO, median preoptic nucleus; MPOL, medial preoptic nucleus, lateral part; SolM, nucleus tractus solitarius; PAG, periaqueductal gray; PVN, paraventricular thalamic nucleus; RMg, raphe magnus nucleus; SFO, subfornical organ; SCh, suprachiasmatic nucleus; VDB, nucleus of the vertical limb of the diagonal band. **G**, Peripheral administrated PTH-biotin binding to the SFO. ***Left,*** Schematic showing peripheral administration of saline, biotin and PTH-biotin to C57 mice. ***Right,*** Immunofluorescence staining of biotin in SFO of C57 mice. Scale bar, 100 μm. **H,** CFos expression in the SFO ***(upper,*** scale bar, 100 μm) and PVN ***(lower,*** scale bar, 100 μm) after peripheral PTH administration. ***Right bar chart,*** Quantification of cFos positive cells in the SFO and PVN (n=3). I, Electrophysiological recording of SFO cells with whole-cell patch (n=8). ***Upper,*** representative electrophysiological whole-cell patch recording of SFO neurons responses to PTH (scale bar, 2μm); ***lower,*** statistical analysis of changes in resting membrane potential during PTH stimulation. RPM, resting membrane potential, P analyzed by unpaired two-tailed t-test. All error bars and shaded areas show mean ± s.e.m. DAPI, 4’,6-diamidino-2-phenylindole, dihydrochloride.

We then performed a PTH-Biotin binding assay to determine whether peripheral administrated PTH can enter the central nervous system and bind directly to brain. Positive PTH-Biotin binding sites were assessed using immunofluorescence staining of biotin, and injection of saline and biotin was also performed as negative controls. The brain nuclei detected with biotin binding includes the SFO, periventricular hypothalamic nucleus (pv), SCh, periventricular hypothalamic nucleus (Pe), AP and the nucleus tractus solitarius (NTS) (Fig. 1G, Supplementary Fig. 2). The majority of identified nuclei were located surrounding the brain ventricles. Jobom et al reported high-level expression of PTH in human cerebrospinal fluid, especially in hyperparathyroidism patients^20^; our results support this by showing that peripheral PTH can indeed enter the central nervous system and bind to specific brain nuclei.

To investigate the brain nuclei responsive to peripheral PTH, we injected human PTH (1-34) or saline intravenously into C57 mice *i.p.* Neuronal activation was assessed following cFos immunostaining, and cFos positive cells were quantified in different brain nuclei. We found that neurons in a variety of brain nuclei, including the SFO, paraventricular thalamic nucleus (PVN), anterior paraventricular thalamic nucleus (PVA) and suprachiasmatic nucleus (SCh) were activated by serum PTH (Supplementary Fig. 3), and the number of cFos positive cells in the SFO and PVN significantly increased after PTH injection (Fig. 1H).

Using the three above-mentioned methodological approaches, we identified the SFO as one prominent brain nucleus that is both physically and functionally connected to the parathyroid gland. The SFO is an important area in the forebrain, a member of the sensory circumventricular organs (sCVO)^19^, which play an important role in the regulation of water and sodium intake^21^. To further confirm the direct effects of PTH on SFO neurons, we used patch clamp recordings to observe the direct neuronal response to PTH. Application of PTH (10^-6^ M) led to significantly increased SFO neuron resting potentials and firing rates (Fig. 1I). In summary, these results demonstrate that the parathyroid glands receive synaptic connections from a variety of brain nuclei, and that SFO neurons directly respond to peripherally administered PTH.

### Activation of SFO^GAD^ and SFO^VGIut^ neurons have different effects on the regulation of PTH and bone remodeling

Neurons in the SFO detect serum nutrients and osmotic pressure directly through leaky blood vessels and regulate drinking and feeding behavior through downstream nuclei^22,26^. To investigate whether neural activity of the SFO modulates serum PTH levels, we activated SFO neurons directly and analyzed serum PTH changes (Fig. 2A). Because calmodulin-dependent protein kinase II (CaMKII) was previously identified as a marker of excitatory neurons in the SFO^22,27^, we activated SFO^CaMKII^ neurons through chemogenetic and optogenetic approach respectively. Chemogenetic modulation was performed by transduction of AAV9-CaMKII-hM3Dq-mCherry into the SFO and Clozapine-N-oxide (CNO) was used to activate SFO^CaMKII^ neurons. We found that acute activation of SFO^CaMKII^ neurons induced 29.0% decrease of serum PTH level 30 mins after CNO administration (Supplementary Fig. 4A). Optogenetic modulation was also performed by transduction of AAV9-CaMKII-ChR2-mCherry into SFO and blue light was used to activate the neurons. Optogenetic stimulation of SFO^CaMKII^ for 15 mins induces 17.1% decrease of serum PTH level (Supplementary Fig. 4B), suggesting a direct response of PTH secretion to SFO neural activities. To study the long-term effect of SFO^CaMKII^ neuron stimulation on the PTH and bone structure, we performed CNO administration every 48 hrs for 1 month. There was significantly lower serum PTH (23.6% lower) in the hM3Dq group than in the control group following activation of SFO^CaMKII^, but no difference in serum calcium levels (Fig. 2A). After chronic stimulation for 4 weeks, bone mineral density (BMD) of the tibiae trabecular bone was 19.6 % lower than the control group (Fig. 2A), accompanied by lower trabecular bone volume fraction (BV/TV) and trabecular number (Tb.N) (Supplementary Fig. 4C). However, the BMD of cortical bone was similar between groups (Fig. 2A). Both the rapid and long-term effect of chemogenetic regulation on PTH level supported a direct causal relationship between SFO activity and PTH regulation.

**Figure 2.**
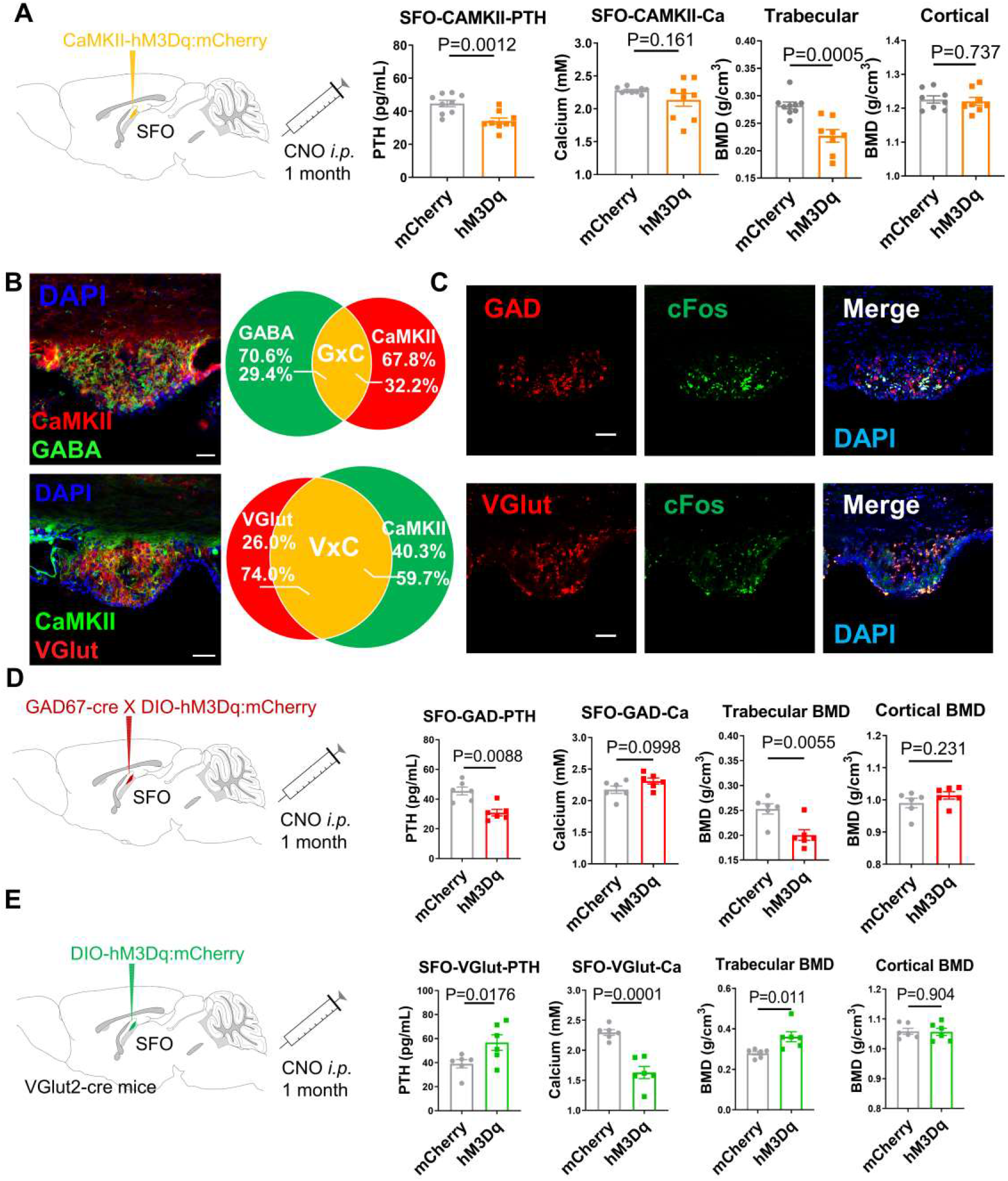
SFO neurons regulate serum PTH level. **A,** Chemogenetic stimulation of Calcium/calmodulin-dependent protein kinase II (CaMKII) neurons in the subfornical organ (SFO) of C57 mice (n=9). ***Left,*** injection of AAV-CaMKII-hM3Dq/mCherry into the SFO of C57 mice followed by 1-month clozapine-N-oxide (CNO) ***i.p.*** administration. ***Right bar charts,*** serum PTH level, total serum calcium level, tibia trabecular bone mineral density (BMD), and femur cortical BMD from MicroCT analysis. **B, *Upper left,*** Immunofluorescence (IF) staining of CaMKII and gamma-aminobutyric acid (GABA) in the SFO of C57 mice. Scale bar, 100 μm. ***Lower left,*** IF staining of CaMKII colocalized with mCherry in the SFO of VGlut2-cre mice transfected with AAV-DIO-mCherry in SFO. Scale bar, 100 μm. ***Upper right,*** statistical analysis of GABA and CaMKII colocalization (n=3 mice). ***Lower right,*** statistical analysis of VGlut and CaMKII colocalization (n=3 mice). **C**, ***In situ*** hybridization of ***Gad1/2*** and ***Vglut2*** mRNAinthe SFO colocalized with IF staining of cFos after ***i.v.*** PTH stimulation in C57 mice. Scale bar, 50 μm. **D**, Chemogenetic stimulation of GABAergic neurons in the SFO of C57 mice (n=6 mice). ***Left,*** injection of AAV-GAD67-cre and AAV-DIO-hM3Dq/mCherry mixture into the SFO of C57 mice followed with 1-month CNO ***i.p.*** administration; ***Right bar charts,*** serum PTH level, total serum calcium level, tibia trabecular BMD, and femur cortical BMD in MicroCT analysis. **E**, Chemogenetic stimulation of glutamatergic neurons in the SFO of VGlut2-cre mice (n=6 mice). ***Left,*** injection of AAV-DIO-hM3Dq/mCherry into the SFO of VGlut2-cre mice followed with 1-month CNO ***i.p.*** administration; ***Right bar charts,*** serum PTH, total serum calcium level, tibia trabecular BMD, femur cortical BMD from MicroCT analysis. P analyzed by unpaired two-tailed t-test. All error bars and shaded areas show mean ± s.e.m. GAD, glutamic acid decarboxylase; VGlut, vesicular glutamate transporter; DAPI, 4’,6-diamidino-2-phenylindole, dihydrochloride.

Previous studies have indicated that the SFO contains intermingled populations of glutamatergic and GABAergic neurons, one which triggers and one which curbs thirst^22^. To explore whether glutamatergic and GABAergic neurons are involved in regulating PTH, we investigated colocalization of CaMKII with gamma-aminobutyric acid (GABA) and vesicular glutamate (VGlut). We found that 32.2% of AAV9-CaMKII-mCherry-labeled neurons expressed GABA in the SFO, whereas 59.7% of CaMKII-labeled neurons colocalize with VGlut (Fig. 2B). We then assessed colocalization of cFos expression and VGlut2/glutamic acid decarboxylase (GAD, enzyme that catalyses the conversion of glutamate to GABA)1&2 expression in SFO induced by peripheral PTH (Fig. 2C). Of the SFO neurons activated following peripheral PTH application, we found an intermingled mixture with 43% glutaminergic and 45% GABAergic neurons (data not shown), suggesting that both type of neurons can be activated by PTH.

To further investigate the regulatory function of GABAergic and glutamatergic neurons in the SFO, we performed chemical genetic stimulation using the AAV9-GAD67-cre virus and the AAV9-DIO-hM3Dq system (Fig. 2D), and we found that specific stimulation of SFO^GAD^ neurons induced a significant decrease of PTH (32.1%) and tibiae trabecular BMD (20.7%), whereas serum calcium levels and cortical BMD were not significantly different from controls (Fig. 2D, Supplementary Fig. 5A). Similarly, we used VGlut2-Cre mice and the AAV9-DIO-hM3Dq system to stimulate SFO^VGlut^ neurons, and chemogenetic stimulation resulted in a 45.3% increase of serum PTH and 29.1% increase in tibia trabecular BMD accompanied by a significant increase in BV/TV, trabecular numbers and trabecular spaces (Fig. 2E, Supplementary Fig. 5B). These results demonstrated that SFO^GAD^ and SFO^VGlut^ neurons both regulate serum PTH levels and bone metabolism, however the effects of these two distinct neuronal populations on the regulation of PTH and bone remodeling appear counter each other, indicating that the CNS bi-directionally regulates PTH level through isolated pathways.

### PTH receptors expressed in SFO neurons regulate negative feedback of PTH secretion

Since peripheral PTH can directly bind and activate SFO neurons (Fig. 1E-F), we next assessed PTH-receptor expression in the SFO and their function in the regulation of serum PTH. In order to study the PTH1R/PTH2R expression in glutamatergic and GABAergic neurons in the SFO, we used colocalization staining and found that PTH1R and PTH2R were both expressed in SFO^VGlut^ and SFO^GAD^ neurons (Fig. 3A-C). Quantification revealed that the majority of cells that expressed PTH1R were GABAergic (81.0%) compared to glutamatergic neurons (53%) (Fig. 3C). Conversely, the majority of PTH2R cells were glutamatergic (73.7%) compared to GABAergic neurons (54.4%) (Fig. 3C).

**Figure 3.**
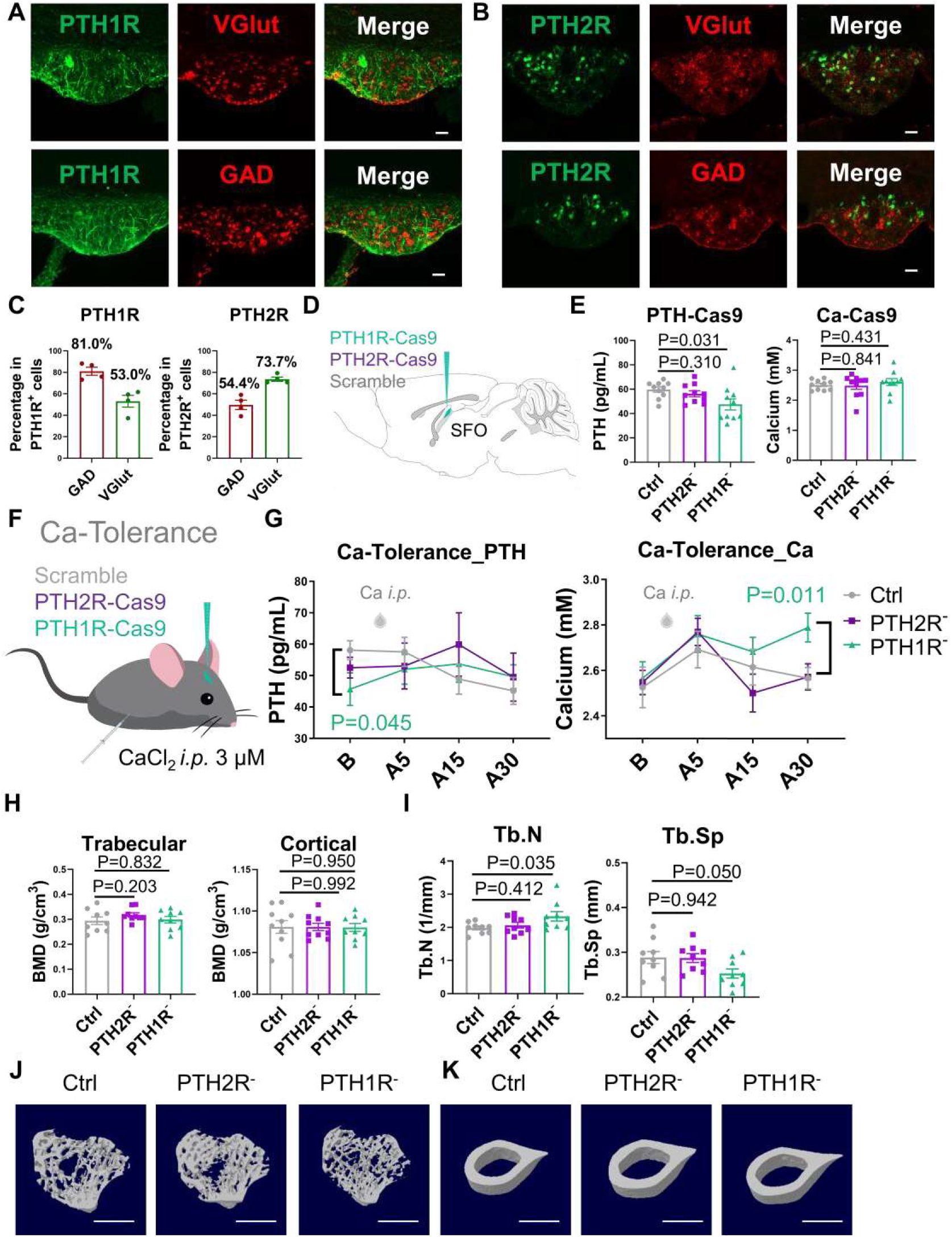
The function of PTH1R and PTH2R in the SFO. **A**, ***Gadl/2*** and ***Vglut2*** mRNA expression colocalized with immunofluorescence staining of PTH1R and PTH2R in the SFO of C57 mice (scale bar, 50 μm). **B**, ***Gadl/2*** and ***Vglut2*** mRNA expression colocalized with PTH2R cells in the SFO of PTH2R-creER^T2^ mice (scale bar, 50 μm). **C**, Statistical analysis of colocalized cells in A and B (n=4 mice, scale bar, 50 μm). **(D-E)** Serum PTH and calcium changes after knockdown of PTH1R/PTH2R in SFO. D, Schematic showing the injection of LV-PTHlR-Cas9 (PTHIR^-^) or LV-PTH2R-Cas9 (PTH2R^-^) or LV-Scramble (Ctrl) into the SFO of C57 mice. **E**, Basic serum PTH and calcium level of C57 mice after knockdown of PTH1R/PTH2R in SFO respectively (n=10 mice). **(F-G),** Calcium tolerance tests in C57 mice after knockdown of PTH1R/PTH2R in the SFO (n=10 mice). **F**, Schematic showing the knockdown of PTH1R/PTH2R, and serum was collected before and 5,15, 30 mins after injection of CaCl_2_ (3μM, *i.p*.). **G**, ***Left***, serum PTH before and after *i.p.* injection of CaCl_2_ in control, SFO^pthlr-^ and SFO^pth2r-^ groups (n=10 mice); ***right,*** serum calcium after ***i.p.*** injection of CaCl_2_ in control, SFO^pthlr-^ and SFO^pth2r-^ groups, (n=10 mice). **H**, Tibia trabecular bone mineral density (BMD) and femur cortical BMD assessed from MicroCT of C57 mice after knockdown of PTH1R/PTH2R in the SFO (n=9 mice). I, Trabecular numbers (Tb.N) and trabecular separation (Tb.Sp) of C57 mice tibia after knockdown of PTH1R/PTH2R (n=9 mice). (J-K), 3D reconstruction of MicroCT analysis of tibia trabecular (J) and femur cortical (K) bones in ctrl/PTH2R’ /PTH1R’ groups (scale bar, 1 mm). P values were analyzed by unpaired two-tailed t-test. All error bars and shaded areas show mean ± s.e.m. VGlut, vesicular glutamate transporter; GAD, glutamate decarboxylase.

Neurons expressing PTH2R are widely distributed in the central nervous system including the locus coeruleus (LC), amygdala, and the periaqueductal gray (PAG), and their functions are correlated with pain and nociceptive behaviors^28–30^. Expression of PTH1R in the SFO was also demonstrated by *in situ* hybridization in the Allen Brain Atlas (Pthlr-RP_110421_01_D07-coronal), however PTH1R function in the brain has not been studied in detail. To investigate the function of the PTH1R and PTH2R in maintaining PTH homeostasis, we down-regulated PTH1R and PTH2R expression in the SFO using LV-PTHlR-Cas9 or LV-PTH2R-Cas9 respectively (Fig. 3D). The knockdown efficiency within SFO was assessed through immunofluorescence staining against PTH1R or PTH2R respectively (Supplementary Fig. 6A-B). The percentage of PTH2R^+^ cells has decreased from 48.3% in control group to 14.5% in PTH2R^-^ group, while the PTH1R^+^ cells has decreased from 53.8% in control group to 23.1% in PTH1R^-^ group (Supplementary Fig. 6C). Serum basal PTH was determined after the knockdown, we found that PTH in PTH1R knockdown group was 20.4% lower than the control group, while PTH level of the PTH2R knockdown group was comparable with the control group (Fig. 3D-E). The serum total calcium remains unaffected in all three groups (Fig. 3E). The effects of knockdown of PTH2R/PTH1R in SFO on bone were also studied by MicroCT analysis. Neither BMD of tibial trabecular bone and femoral cortical bone was affected, however Tb.N increased and Tb.Sp decreased after the knockdown of PTH1R compared to the control group (Fig. 3H-K, Supplementary Fig. 6D).

To further investigate the effects of the parathyroid hormone receptors on maintaining PTH homeostasis, animal was administrated with 3 μM of CaCl_2_ *i.p.* to induce hypercalcemia and the serum PTH and total calcium was assessed before and 5,15, 30 min after the stimulation (Fig. 3F-G). In response to acute calcium increase, the serum PTH of control group show consistent decrease from the 15^th^ min (Al5) to 30^th^ min after injection (A30). The serum calcium level of the control group increased by A5 and decreased rapidly afterwards and returned to normal by the time A30. However, for the SFO^PTH1R-^ and SFO^PTH2R-^ groups, both PTH level increased after calcium stimulation and reached peak by A15 and began to decrease indicating a later latency of PTH regulation response. The serum calcium level of the SFO^PTH2R-^ group increase and reversed by the same time of control group with larger amplitude than the control group, however, the calcium level of SFO^PTH1R-^ group failed to return to normal by A30 (Fig. 3G). These results indicate that knockdown of PTH receptors in SFO not only affected PTH basal secretion, but also affect the kinetics of PTH in response to hypercalcemia stimulation.

We also performed the fluid consumption tests in the SFO^PTH1R^ and SFO^PTH2R^ knockdown mice since SFO is important organ for water-salt balance. It was found knockdown of PTH1R in SFO induced a decreased water consumption compared to control group, while PTH2R knock down in SFO did not show an effect (Supplementary Fig. 6E).

Previous studies of the function of PTH2R in brain were mainly performed through universal knockout experiments^30,31^. To further gain more insight into the function of PTH2R neurons in SFO, we constructed a PTH2R-creER^T2^ transgenic mouse strain (Supplementary Fig. 7–9). The successful construction of a PTH2R-creER^T2^ mice was validated though cross breading of PTH2R-creER^T2^ mice with Ai14 mice resulting in tdTomato protein expression in Cre-recombinase expressing cells. Immunofluorescence staining of PTH2R was performed in PTH2R-creER^T2^ X Ail4 mice brain, and positive signals was observed in SFO, Arc, PVN, MPOL and a variety of brain nuclei previously reported with high PTH2R expression^32^ (Supplementary Fig. 7A, Supplementary Fig. 9). Colocalization staining with NeuN also demonstrated that the majority of PTH2R^+^ cells in the SFO are neurons (Supplementary Fig. 7B). Then we investigated the function of PTH2R expressing neurons in SFO, chemogenetic activation of PTH2R neurons resulted in a 20.9% increase in serum PTH and no changes in serum calcium (Supplementary Fig.7E), which is consistent with the effects of activation of SFO^VGlut^ neurons. However, trabecular and cortical BMD was not affected (Supplementary Fig. 7F). In the open field study, mice in hM3Dq group spend less time in central areas while central entries, and total distance was not affected (Supplementary Fig. 8A). There was also no difference between groups in the total immobility time in tail suspension test (Supplementary Fig. 8B). Chemogenetic activation of SFO^PTH2R^ also led to a higher intake of water compared to the control group (Supplementary Fig. 8C). Taken together, our data demonstrated that PTHR expressing neurons in SFO are essential in both sensing PTH/calcium levels and regulating PTH secretion; PTH1R and PTH2R play different roles in regulating the basal PTH secretion, PTH homeostasis in response to exogenous CaCl_2_ stimulations and bone remodeling.

### PVN regulation of PTH

Next, we investigated the downstream effectors of SFO neurons. We know that SFQ^CaMKII^ neurons project to a variety of brain nuclei including the MnPO, paraventricular nucleus (PVN), Vascular organ of lamina terminalis (OVLT) and bed nucleus of the stria terminalis (BNST)^21^ (Supplementary Fig. 10). Of these brain nuclei, the PVN is reported as crucial in the regulation of both neuroendocrine homeostasis and sympathetic activity, then we investigated whether PVN neurons are involved in the regulation of PTH and bone metabolism.

Unlike the SFO, which consists of both GABAergic and glutamatergic neurons, we found that the majority of PVN neurons responding to peripheral PTH stimulation (cFos-positive) were colocalized with VGlut2 and not GAD1&2 (Fig. 4A). Immunofluorescence showed that CaMKII and VGlut2 are both abundantly expressed in the PVN. Quantification revealed that VGlut2/CaMKII double-positive cells accounted for 62.6% of VGlut2-positive neurons, and 67.7% of CaMKII-positive neurons (Fig. 4B).

**Figure 4.**
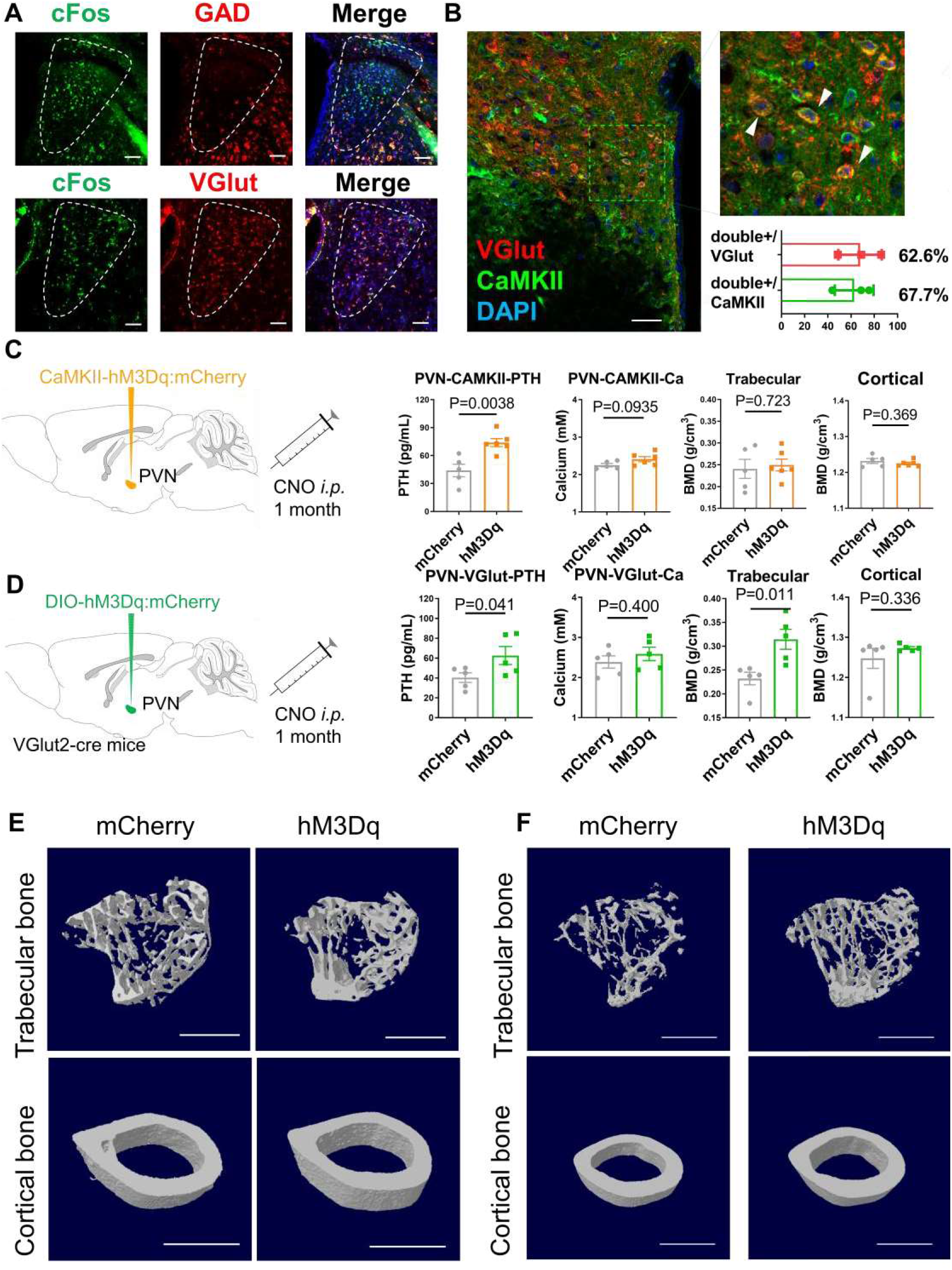
PVN neurons regulate serum PTH level. **A**, ***In situ*** hybridization of GAD 1/2 and VGlut2 mRNA in the paraventricular nucleus (PVN) colocalized with cFos staining after *i.v.* PTH stimulation. Scale bar, 100 μm. **B**, ***Left and upper right,*** colocalization of VGlut and CaMKII markers in the PVN of VGlut2-cre mice (scale bar, 100 μm); ***lower right***, statistical analysis of the colocalized cells (n=3 mice). **C**, Chemogenetic activation of CaMKII neurons in the PVN of C57 mice (n=5-6 mice). ***Left,*** schematic showing the injection of AAV-CaMKII-hM3Dq/mCherry into the PVN of C57 mice followed with 1-month CNO *i.p.* administration. ***Right bar charts,*** serum PTH, total serum calcium level, tibia trabecular bone mineral density (BMD), and femur cortical BMD from MicroCT analysis. **D,** Chemogenetic stimulation of VGlut neurons in the PVN of VGlut2-cre mice (n=6 mice). ***Left,*** schematic showing the injection of AAV-DIO-hM3Dq/mCherry into the PVN of VGlut2-cre mice followed with 1-month CNO *i.p.* administration; ***Right bar charts,*** serum PTH, total serum calcium level, tibia trabecular BMD, and femur cortical BMD from MicroCT analysis. **E**, 3D reconstruction of MicroCT analysis of tibia trabecular bone and femur cortical bone in C57 mice transfected with CaMKII-mCherry or CaMKII-hM3Dq-mCherry in the PVN and then administrated with CNO for 1 month (scale bar, 1 mm). **F**, 3D reconstruction of MicroCT result of tibia trabecular bone and femur cortical bone in VGlut2-cre mice transfected with DIO-hM3Dq-mCherry or DIO-mCherry in PVN and administrated with CNO for 1 month. Scale bar, 1 mm. P analyzed by unpaired two-tailed t-test. All error bars and shaded areas show mean ± s.e.m. CNO, clozapine-N-oxide.

Next, we investigated the regulatory functions of pvN^CaMKII^ and PVN^VGlut^ neurons. Chemogenetic activation of pvN^CaMKII^ neurons induces 63.0% increase in serum PTH level 30 min after CNO administration (Supplementary Fig. 11A-B). After chronic chemogenetic activation of PVN^CaMKII^ neurons for 1 month, the serum PTH level in the hM3Dq group became 62.2% higher than the mCherry group (Fig. 4C). There was no difference in the serum total calcium, BMD of tibial trabecular or femoral cortical bone or trabecular microstructures between groups (Fig. 4C,4E, Supplementary Fig. 11C). Interestingly, we activated PVN^VGlut^ neurons after transduction hM3Dq through injection of the AAV9-DIO-hM3Dq-mCherry into the PVN of VGlut2-Cre mice (Fig. 4D) and found that chronic activation of PVN^VGlut^ neurons led to significantly higher serum PTH than the control group (55.0%), and a higher trabecular BMD in mice tibiae (35.4%) (Fig. 4D, Fig. 4F). Both percent bone volume and trabecular separation were significantly changed in the experimental group (Supplementary Fig. 11F). These data suggest that the activation of glutamatergic neurons in PVN efficiently up-regulate PTH levels and bone remodeling.

### GABAergic neurons in the SFO send neuronal projections to the PVN and regulate bone remodeling

Direct SFO^CaMKII^ projection to the PVN has been observed by Zimmerman, et al. ^25^. However, it is not known whether the PVN receives an SFO^GAD^ projection. We studied SFO^GAD^ neuronal projections using a combined injection of AAV9-DIO-mCherry and AAV9-GAD67-Cre viruses into the SFO of C57 mice. In addition to the previously described SFO^VGAT^→MnPO and SFO^VGAT^→OVLT neuronal projection (VGAT, vesicular GABA transporter, that loads GABA from the neuronal cytoplasm into synaptic vesicles)^22,26^, we found that the PVN also receives dense innervation from SFO^GAD^ neurons (Fig. 5A). To confirm that the PVN^VGlut^ neurons specifically receive synaptic connections from SFO^GAD^ neurons, we performed retrograde tracing by injecting AAV9-DIO-TVA-RVG and RV-ΔG-mCherry into the PVN of VGlut2-Cre mice (Fig. 5B). Efficient transfection of AAV helper and RV in PVN glutamatergic neurons was indicated by mCherry signals in the PVN. In the SFO, RV positive mCherry signals, which were colocalized with GABA and CaMKII, indicated retrograde monosynaptic connection with PVN^VGlut^ neurons (Fig. 5B).

**Figure 5.**
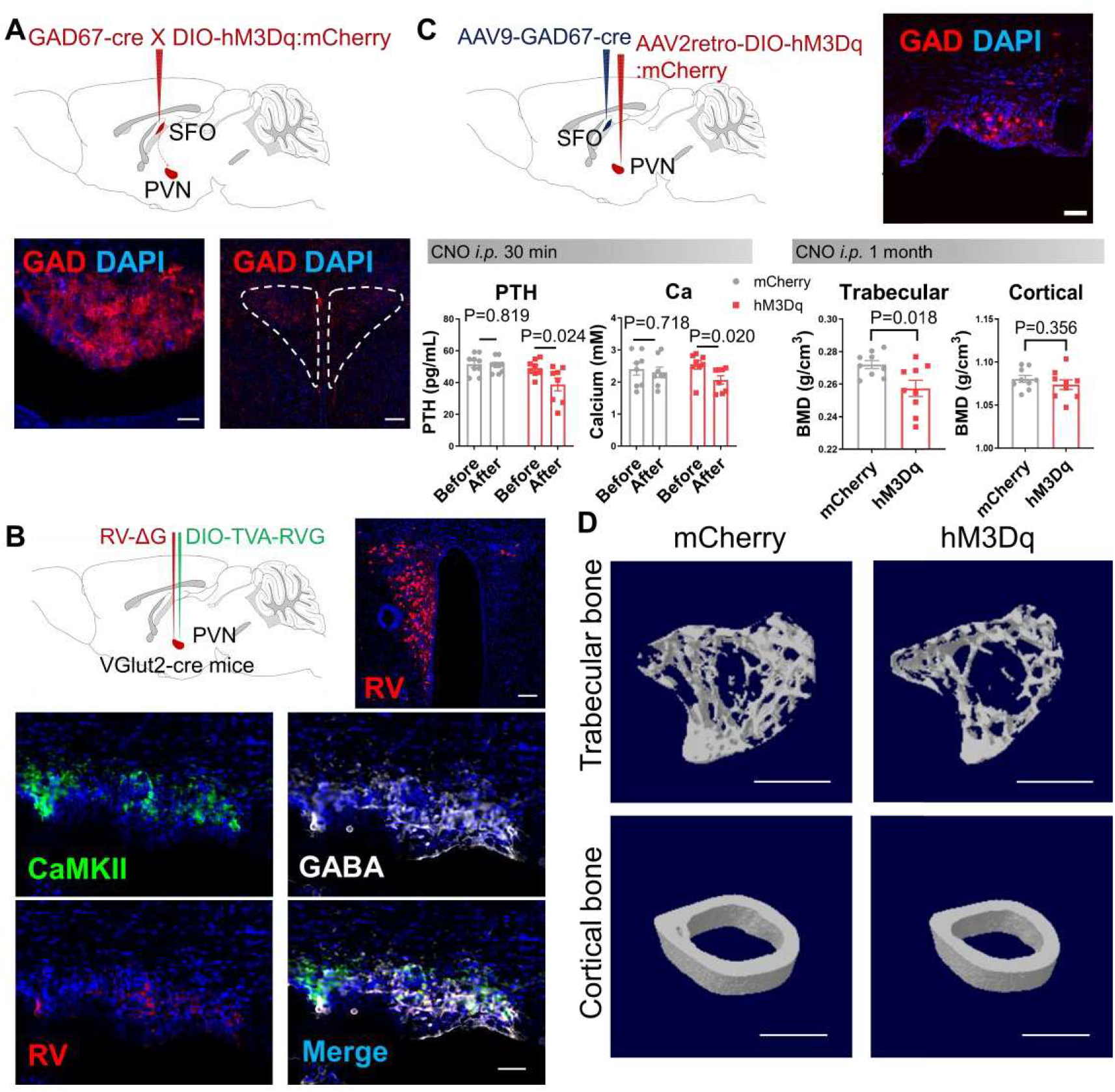
SFO^GAD^ neurons send projections to the PVN. **A**, ***Upper,*** schematic showing the injection of AAV-GAD67-cre and AAV-DIO-mCherry mixture into the SFO of C57 mice; ***lower,*** GABAergic neural projections observed from the SFO ***(left,*** scale bar, 50 μm) to the PVN ***(right,*** scale bar, 100 μm). **B**, ***Upper left,*** schematic showing injection of AAV9-DIO-TVA-RVG and then RV-ΔG-dsRed into the PVN of VGlut2-cre mice; ***upper right,*** genetically modified rabies virus (RV) tracing from PVN^VGlut^ neurons (scale bar, 100 μm). ***Middle and lower,*** colocalization of RV signals with IF staining of CaMKII and GABA signals in SFO (scale bar, 50 μm). **(C-D)** Chemogenetic activation of SFO^GAD^-→PVN neural projections in C57 mice (n=9 mice). **C, *Upper left,*** schematic showing injection of AAV-GAD67-cre into the SFO and AAV2retro-DIO-hM3Dq/mCherry into the PVN of C57 mice followed with 1-month CNO ***i.p.*** administration. ***Upper right,*** mCherry signals observed in the SFO indicating GAD67^+^ neurons with axon terminals projected to PVN (scale bar, 50 μm). ***Lower left,*** serum PTH before and 30 min after CNO injection, total serum calcium level before and 30 min after *i.p.* CNO injection into mCherry group and hM3Dq group. ***Lower right,*** tibia trabecular BMD and femur cortical BMD in MicroCT analysis after 1-month CNO administration into mCherry group and hM3Dq group. **D,** 3D reconstruction of MicroCT analysis of tibia trabecular bone and femur cortical bone in C57 mice injected with GAD67-cre virus in the SFO and DIO-mCherry or DIO-hM3Dq-mCherry in the PVΓ✓ and then given CNO for 1 month. Scale bar, 1 mm. P analyzed by paired two-tailed t-test for before-after analysis, unpaired two-tailed t-test for mCherry-hM3Dq analysis. All error bars and shaded areas show mean ± s.e.m.

To explore regulatory function in SFO^GAD^-PVN circuit, we injected AAV9-GAD67-Cre virus into the SFO and AAV2retro-DIO-hM3Dq-mCherry virus into the PVN of C57 mice. Specific chemogenetic activation of the neural projections from SFO^GAD^ – PVN circuit induced 20.8% decrease of serum PTH level and 18.4% decrease of serum calcium level within the hM3Dq group (Fig. 5C). The tibial trabecular BMD was also decreased by 5% while femoral cortical BMD was not affected (Fig 5C-D). Trabecular BV/TV and Tb.N were also decreased in hM3Dq group (Supplementary Fig. 12A). In the behavior test, the water and salt consumption were not affected during the activation of the SFO^GAD^→PVN circuit (Supplementary Fig. 14B). These results reveal a novel SFO^GAD^-PVN circuit which specifically regulates serum PTH, total calcium and trabecular bone metabolism.

### Ablation of sympathetic nerves affect serum PTH level

Innervation of parathyroid glands has been reported in previously studies. In our study, we observed both the sympathetic nerves (TH^+^) and sensory nerves which were labeled with calcitonin gene related protein (CGRP^+^) in the parathyroid gland (Fig. 6A-B). Double staining revealed sympathetic nerves in mice parathyroid glands, which form a net-like structure that surrounds the blood vessels (CD31^+^); the sympathetic nerves form clusters with blood vessels rather than an even distribution in the parathyroid glands (Fig. 6B). On the other hand, most CGRP nerves were observed between parathyroid-gland substructures as single neural fibers (Fig. 6B).

**Figure 6.**
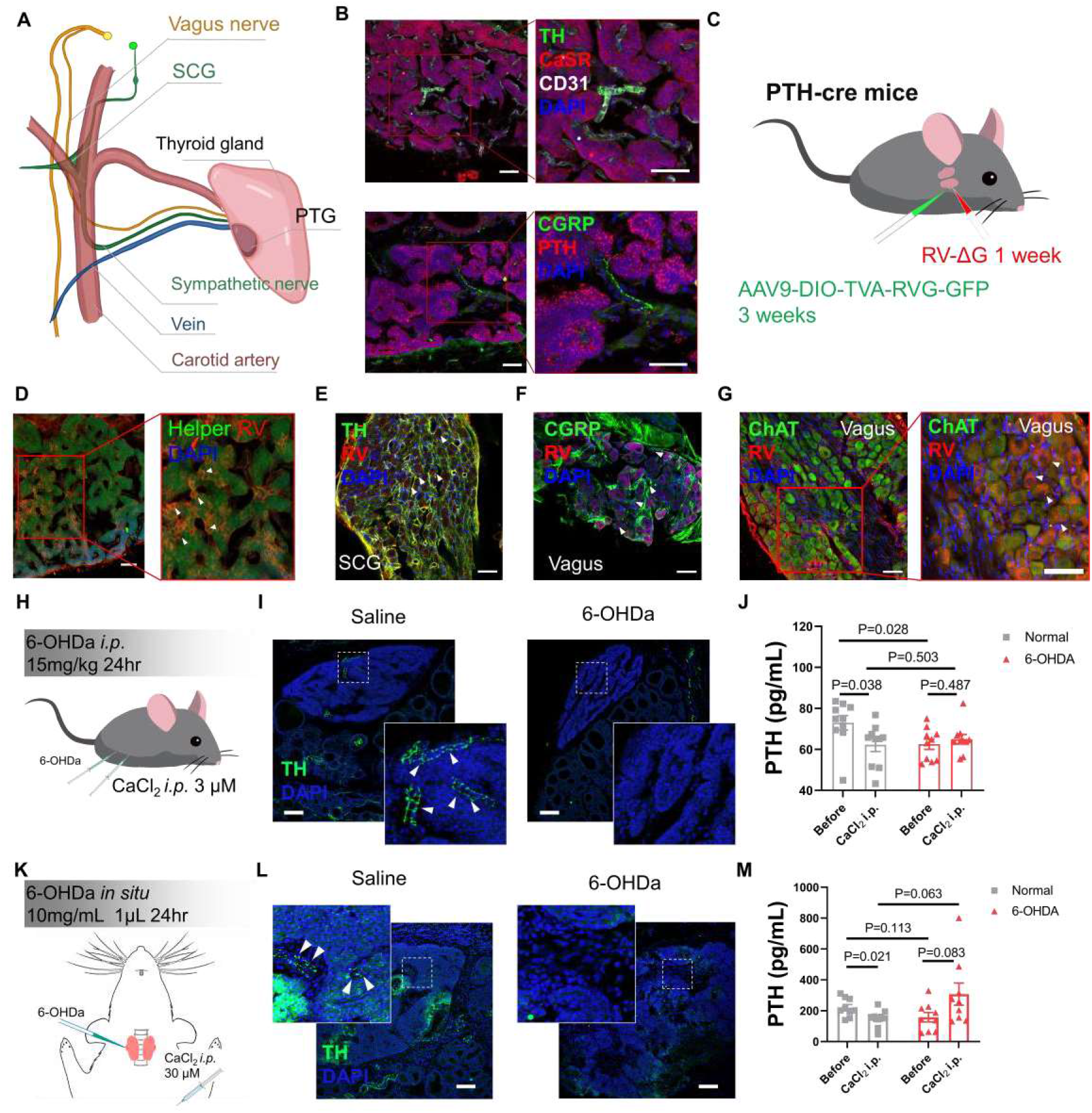
Peripheral innervation in parathyroid glands and sympathetic regulation of PTH. **A,** Schematic demonstration of the anatomical relationship between parathyroid gland (PTG) and the surrounding peripheral nerves and blood vessels. PTG, parathyroid gland; SCG, superior cervical ganglia. **B**, Innervation and location of tyrosine hydroxylase (TH) sympathetic nerves, CD31 capillaries and calcitonin gene-related peptide (CGRP) sensory nerves in mice parathyroid glands. Scale bar, 50 μm. **C**, Schematic showing helper (AAV-DIO-TVA-RVG-GFP) and rabies virus (RV-ΔG-dsRed) infection of the parathyroid gland in PTH-cre mice and trans-monosynaptic connection to the peripheral nervous system. **(D-G),** Neural specific rabies virus (RV) tracing from mice parathyroid glands to the superior cervical ganglion (SCG) and the inferior ganglion of vagus nerve (Vagus). **D, *In situ*** RV (red) and helper (green) expression in the parathyroid gland of PTH-cre mice. White arrow indicates colocalization of RV expression and helper signals. Scale bar, 50 μm. **E**, RV signals colocalized with IF staining of TH expression in mice SCG. **F,** RV signals colocalized with IF staining of CGRP expression in the vagus ganglion. **G,** RV signals colocalized with IF staining of ChAT expression in the vagus ganglion. White arrow indicates colocalization of RV signals with TH/ChAT/CGRP signals. Scale bar, 50 μm. **(H-J),** Calcium tolerance study in C57 mice of universal ablation of sympathetic nerves. H, Schematic showing administration of 6-OHDa (15 mg/kg, *i.p*.) to ablate the sympathetic terminals and calcium tolerance test was performed 24hrs after 6-OHDa injection. **I,** IF staining of TH indicates that sympathetic nerve innervation in mice PTG were ablated 24hrs after 6-OHDa administration. Scale bar, 100 μm. **J**, Serum PTH levels before and 30 mins after *i.p.* administration of CaCl_2_ in C57 mice with or without sympathetic nerve ablation by 6-OHDA (n=lO mice). P analyzed by two-way ANOVA test comparing the 6-OHDa effects among different groups. (K-M), Calcium tolerance study in SD rat of PTG after local ablation of sympathetic nerves. **K**, Schematic showing local injection of 6-OHDa (10 mg/mL, lμL) to ablate the sympathetic nerve in rat parathyroid glands and then calcium tolerance test was performed 24hrs after 6-OHDa injection. **L**, IF staining of TH indicating sympathetic nerve innervation in rat PTG were ablated 24hrs after 6-OHDa administration. Scale bar, 100 μm M, Serum PTH before and 30 mins after *i.p.* administration of CaCl_2_ in rats with or without sympathetic nerve ablation by 6-OHDA (n=9 rats). The P value were analyzed by unpaired two-tailed t-test. All error bars and shaded areas show mean ± s.e.m. 6-OHDA, 6-hydroxydopamine; CaSR, calcium sensory receptor; CD3l, cluster of differentiation 31; CGRP, calcitonin-gene related protein; ChAT, choline acetyltranserase; PTG, parathyroid gland; SCG, superior cervical ganglia; TH, tyrosine hydroxylase.

Physical connectivity between the parathyroid glands and the nervous system was also investigated using neuronal-specific retrograde tracing from the parathyroid glands. We injected CTB-Alexa Fluor 488 into one side of the rat parathyroid glands and then after sacrifice, both side of the sympathetic superior cervical ganglia (SCG) and inferior vagal ganglia (Vagus) were isolated to study innervation. CTB signals were observed in both sides of the SCG and Vagus (Supplementary Fig. 13). These results are consistent with the findings of both sympathetic nerves and CGRP nerves in parathyroid glands.

Next, we performed RV tracing using PTH-Cre mice to assess whether there is direct synaptic connectivity between parathyroid chief cells and the peripheral nervous system. The signals of both RV and helpers were observed in parathyroid glands in PTH-Cre mice, indicating that the AAV helper and RV had infected parathyroid chief cells (Fig. 6C-D). One week after virus infusions, positive RV signals were observed in the SCG and Vagus following counterstaining of tyrosine hydroxylase (TH) in the SCG, choline acetyltransferase (ChAT) and calcitonin gene-related protein (CGRP) in the Vagus (Fig 6E-G). These results indicate that the SCG and Vagus may be directly connected to the parathyroid chief cells by direct, monosynaptic connections.

The function of peripheral sympathetic nerves in the regulation of PTH was investigated further through ablation of peripheral sympathetic neural terminals by 6-OH-Dopamine. The sympathetic nerve terminals were ablated universally in mice through *i.p.* administration in mice (Fig. 6H-J) and locally within the parathyroid glands in rats (Fig. 6K-M). Universal ablation of sympathetic neural terminals in mice induced a decrease of basic serum PTH (Fig. 6J), and the serum PTH level was also not responding to the hypercalcemia stimulation (Fig. 6J). After the ablation of the sympathetic nerve terminals in PTG of rats, basal PTH level were not affected, however the serum PTH was not responding to the hypercalcemia stimulation in the 6-0HDa group compared with the control group (Fig. 6M). Taken together, these data consistently demonstrate that the sympathetic and sensory nervous system are both tightly integrated with the parathyroid glands, and that the ablation of peripheral sympathetic nerve significantly impacts PTH response to hypercalcemia stimulation.

## Discussion

Modulation of serum calcium levels and bone remodeling relies upon serum PTH, the most important hormone in this regard, levels of which are maintained through humoral, hormonal and neural regulation^33^. However, until now the effects of central neural regulation of serum PTH and underlying mechanisms were largely unknown. Using neural circuit tracing and chemogenetic techniques in addition to analysis of PTH and bone metabolism, we for the first time, systemically studied the structural and functional connection between the parathyroid and specific types of neurons in the central nervous system. This resulted in the identification of the components of a specific central neural circuit that modulates PTH and in the revelation of the underlying mechanism of central neural regulation of PTH levels and bone metabolism (Supplementary Fig. 14).

### Central nervous system (CNS), PTH and bone metabolism

The close relationship of PTH to the CNS has been observed for decades. The earliest studies suggesting PTH function within the CNS stem from a series of findings beginning with the identification of PTH-like immunoreactive protein in the brains of different species, particularly in bony fish. In mammals, the embryonic origin of parathyroid gland is derived from the 3^rd^ and 4^th^ brachial pouch, which closely neighbors the neural crest (NC) mesenchyme^34,35^. An evolutionary model has also been proposed, in which, during the evolutionary period where aquatic vertebrates transitioned to a terrestrial environment, the parathyroid gland gradually migrated from the CNS to the periphery to enhance calcium regulatory capacity during times of unstable calcium dietary supplementation^2^. However, the question of whether the CNS still reserves calcium and PTH regulation functionality in adult mammals has never been asked. In this study, we provide the first demonstration that chemogenetic activation of SFO^GAD^, SFO^VGlut^, PVN^CaMKII^ and PVN^VGlut^ neurons are able to directly modulate serum PTH levels and the bone metabolism in the absence of any other experimental manipulation. Conversely, knocking down the expression of PTH1R and PTH2R in the SFO, or ablation of the sympathetic nerves in the PTG, led to a significant blockade of serum PTH and calcium responses to exogenous calcium stimulation. This evidence demonstrates the necessity of the centred and peripheral nervous systems in both regulating PTH basal secretion and maintaining PTH homeostasis through feedback loops.

Parathyroid hormone is an important hormone in the bone remodeling process, where it initiates bone resorption followed by bone formation^1^. Ablation of PTH4 neurons result in abnormal bone mineralization and significantly affects osteoblast differentiation^8^, suggesting that PTH might be an important regulator of the central neural control of bone remodeling. In our study, we found that changes in bone mass were typically in the same direction as PTH changes; that is, decreased PTH was followed by decreased tibiae trabecular BMD and increased PTH was followed by increased tibiae trabecular BMD. Previous studies have indicated that intermittent administration (daily) of PTH induces an anabolic effect in bone, which is contrary to the effects of continuous administration^1^. In our study, chemogenetic activation of brain neurons was performed with transduction of hM3Dq in the brain and administration of CNO (*i.p*.) every 48 hours. Based on a pharmacokinetic study of CNO^36^, serum PTH changes induced by chemogenetic stimulation of mice reached peak levels every 48 hours, which may explain the anabolic effects of PTH on bone.

### SFO neurons are important in both the detection and regulation of PTH level

Both the short-term and long-term effect of chemogenetic regulation on PTH level supported the direct causal relationship between SFO activity and PTH regulation. As a member of the CVO, the SFO directly detects circulating molecular signals in serum.

We found PTH1R and PTH2R expression in SFO neurons, reaffirming this finding reported in the Allen Brain Atlas database. The binding efficiency of PTH1R to PTH is different from that of PTH2R. The EC5O of PTH1R to PTH (1-34) is around 1-2 nM, whereas the EC5O of PTH2R to PTH (1-34) is around 70 nM^37^. Therefore, serum PTH in healthy individuals (65-150 pg/mL) would likely predominantly activate only PTH1R in the SFO. In our study, we found that PTH1R expression was predominant in SFO^GAD^ neurons and activation of SFO^GAD^ neurons induced a decrease in serum PTH. It is thus expected that, under physiological conditions, the CNS can modulate serum PTH in a negative feedback pathway mainly through SFO^GAD^ neurons.

On the other hand, under severe pathological conditions such as hyperparathyroidism (HPT), serum PTH levels are around 10-100 times higher than physiological levels and SFO^VGlut^ neurons, which mainly expressed PTH2R, would be activated to induce a consistent increase of serum PTH. Our study indeed found that chemogenetic activation of the SFO^VGlut^ neurons did induce a significant elevation of PTH levels and increased bone density, which is a reverse of the regulatory effects of SFO^GAD^ neurons. Our findings may thus explain why serum PTH becomes uncontrollable in some patients of late-stage HPT and leaves them with no other treatment except parathyroidectomy surgery.

In addition, it is consistently reported that HPT patients develop psychiatric symptoms including anxiety, depression and cognitive disorder^38^. Moreover, if these symptoms develop in primary or secondary HPT diseases, they improve following parathyroidectomy surgery^17,39–41^. The function of PTH2R and the agonist TIP39 mediate nociception, including neuropathic and inflammatory pain^29^; genetic knockdown of PTH2R in the medial amygdala and disruption of PTH2R projections to the bed nucleus of the stria terminalis (BNST) influence fear- and anxiety-like behavior ^30,42^. However, there Eire no gain of function studies for PTH2R reported in the literature. In our study, we firstly found that knockdown of *pth2r* in the SFO did not induce behavioral changes. However, chemogenetic activation of SFO^PTH2R^ neurons in PTH2R-creER^T2^ mice led to less time in the center during OFT tests, which is used as an index of anxiety, even although immobility time during TST was not affected. These data indicate that PTH2R neurons in the SFO regulate anxiety-like behaviors in mice, in addition to regulating PTH level and the bone remodeling process.

### The PVN, sympathetic nerves and the PTH

Of the nuclei downstream of the SFO, we chose the PVN as the target to study its function in the regulation of PTH. In addition to the fact that PVN receives polysynaptic connections from the PTG and is responsive to peripherally administrated PTH and calcium, the PVN receives a direct monosynaptic neural projection from SFO GABAergic neurons as revealed by our RV tracing study. The PVN is well known as crucial member of the neural-humoral hypothalamic-pituitary-thyroid (HPT) axis and the hypothalamic-pituitary-adrenal (HPA) axis^43^. In addition, PVN^VGlut^ neurons also directly project to the nucleus tractus solitarius (NTS), which affects sympathetic nerves through a neural sympathetic connection^44^; activation of PVN^VGlut^ neurons increase the activity of the sympathetic nervous system^45^.

By directly activating PVN neurons, we primarily found that activation of PVN neurons results in elevated serum PTH levels, and found different regulatory effects following activation of PVN^CaMKII^ or PVN^VGlut^ neurons, despite overlap of these neurons. Activation of both pvN^CaMKII^ and PVN^VGlut^ induced an increase in serum PTH, however, trabecular BMD was not affected following pvN^CaMKII^ activation yet increased following PVN^VGlut^ activation. These findings suggest that PVN^VGlut^ neurons are more likely to regulate both PTH levels and bone metabolism compared to pvN^CaMKII^ neurons, since PVN^VGlut^ neurons directly receive a monosynaptic projection from SFO^GAD^ neurons. Previous research has indicated that noradrenergic neurons in the PVN suppress stress-induced bone loss^46^, whereas our study shows that PVN^VGlut^ neurons exerted more of an effect on bone metabolism through the regulation of PTH. The inhibitory effects on trabecular bone BMD following specific activation of the GABAergic neural projection from the SFO to the PVN further confirmed that PVN can relay upstream signals from the SFO to regulate bone metabolism (Figure 5). However, neural activation of SFO and PVN may also lead to systematic effects on other organs resulting in indirectly influence on PTH and bone, which requires further study in future.

Previous work has indicated that serum PTH increased tenfold during electric stimulation of the cerebral sympathetic trunk (CST)^16^. Consistent with this, we found that ablation of peripheral sympathetic nerves in mice induced a significant decrease of serum PTH. This finding is also consistent with an earlier study in which direct infusion of epinephrine induced 127 to 255% increase of PTH in cows^15^. From our immunofluorescence study, we found that TH nerve terminals were tightly surrounded by the capillaries within the PTG, indicating that serum PTH may be regulated sympathetically via capillary vasodilation and contraction. The neural tracing experiments and ablation study further confirmed that sympathetic innervation is necessary for relaying the central signals to regulate PTH homeostasis.

In summary, our contributions are in three parts. Firstly, we identified the SFO and PVN as important nodes that regulate serum PTH levels and bone metabolism. Secondly, for the first time, we revealed a population of GABAergic neurons in the SFO, which not only detect PTH levels via PTH receptors, but also directly regulates PTH levels. Thirdly, we found that the PVN and peripheral sympathetic system were downstream of the SFO in this circuit and play important roles in the regulation of PTH homeostasis and bone metabolism. Therefore, not only did we identify the underlying mechanism of central neural regulation of serum PTH, but also provide evidence of the intricate process by which the CNS detects peripheral PTH and the responses to changes in serum PTH under physiological or pathological conditions.

## Methods

### Animals

C57/6 J mice and SD rats (Beijing Vital River Laboratory Animal Technology Co) raised at the Shenzhen Institute of Advanced Technology, Chinese Academy of Science were used. VGlut2-ires-cre mice (Jackson Laboratory, 016863), PTH-cre mice (Jackson Laboratory, 005989), Ail4 mice (Jackson Laboratory, 007908) were also used. PTH2R-CreER^T2^ mice were established and provided by GermPharmatech Co., Ltd. Only male mice/rats of 6-8 weeks-old were used in this study. All animals were housed under a Specific Pathogen Free (SPF) barrier environment according to the standard of GB14925-2010 Laboratory Animals Requirement of Housing and Facilities within a temperature-controlled room (20-26°C) with a 12-hour light/dark cycle (07:00 lights on, 19:00 lights off). Animals had free access to chow and water. All studies and experimental procedures were approved by the Institutional Animal Care and Use Committee (IACUC) of the Shenzhen Institute of Advanced Technology, Chinese Academy of Science.

PTH2R-CreER^T2^ mice were generated based on C57/6 J mice using the CRISPR-Cas9 technique by inserting a strand of ere recombinase with an estradiol receptor (ERT2) after the PTH2r promoter. Tamoxifen (75 mg/kg, i.p.) was administrated to the animal 5 days before experiments to initiate ere recombinase gene expression.

### Retrograde neural tracing from parathyroid glands

Neural specific retrograde tracing was conducted with chloral toxin B (CTB-Alexa Fluor 488,10 mg/mL, Molecular Probes^®^), modulated pseudorabies virus vector (PRV), modulated rabies virus vector (RV) and its adeno-associated virus (AAV) helper. Non-trans-synaptic retrograde tracing was conducted by CTB, which is absorbed by the neural terminals and can be transported to the neuronal soma. Monosynaptic retrograde tracing was performed with sequential use of AAV helper (DIO-EGFP-TVA-RVG) and modulated RV (RV-EnvA-ΔG-dsRed). The AAV helper was used for Cre-dependent expression of TVA, which directs the infection of RV and G proteins that reinstate the trans-synaptic infection. Poly-synaptic retrograde tracing was employed by pseudorabies virus (PRV-GFP), which infects the neural terminals of the injection site and is delivered to the soma and transfects to the next neuron through synaptic connection.

Helper AAV, RV and PRV (BrainVTA, Wuhan, China) were used with titers of 1.3E+12 PFU/mL, 5.0E+08 PFU/mL and 2.0E+09 PFU/mL, respectively. The retrograde tracers were injected into the parathyroid glands of mice and rats using a Hamilton^®^ microsyringe under anaesthesia (*i.p.* 100mg/kg sodium pentobarbital). The precise injection into mice PTG was performed through microscopic surgery and exclusion of animals were carried out for the off-target injections.

### Virus injection and chemogenetic regulation of neurons

Adult mice of 6-8 weeks were anaesthetized (*i.p.* 100mg/kg sodium pentobarbital) and mounted on a stereotaxic frame (RWD Life Science, Shenzhen, China). Virus suspension (AAV9-CaMKII-mCherry, AAV9-CaMKII-hM3Dq-mCherry, AAV9-CaMKII-ChR2-mCherry, AAV9-DIO-mCherry, AAV9-DIO-hM3Dq-mCherry, AAV9-GAD67-cre, AAV2retro-DIO-hM3Dq-mCherry, AAV2retro-DIO-EYFP, LV-PTH1R-Cas9, LV-PTH2R-Cas9, LV-empty; 1.5-2.5E+12 PFU/mL) were loaded into a 5μl-Hamilton^®^ syringe with 33-g needle, and 150 nL was injected with a speed of 100 nL/min by the targeting locations (SFO: −0.58, 0, −2.5; PVN: −0.8, ±0.5, −4.5) based on a stereotaxic atlas^47^. The surgical procedure was based on Lowery and Majewska^48^. After surgery, animals were allowed to recover for at least 1 week before experiments. In chemogenetic studies, CNO (*i.p.* 1 mg/kg, MCE, 34233-69-7) was injected 3 weeks after virus expression, every 48 hours for 4 weeks prior to serum and bone collection and analysis. The AAV9 virus were all prepared in our lab, and the LV-PTHlR-Cas9, LV-PTH2R-Cas9 and LV-empty were accessed from BrainVTA, Wuhan, China. The precise injection into mice brain was performed and exclusion of animals were carried out for the off-target injections.

### Optogenetic stimulation of CaMKII neurons in SFO

C57 mice were injected with AAV9-CaMKII-ChR2-mCherry or AAV9-CaMKII-mCherry into the SFO through stereotaxic injection. Three weeks after virus injection, an optical fiber (200 μm diameter) was implanted 0.2 mm above with SFO with an cannula fixed onto the skull. Optogenetic stimulation was performed 1 week after optic fiber implantation. The light stimulation was performed with 450 nm blue light and stimulated under 0.5-1 mW/cm^2^, 20 Hz for 15 min.

### PTH and calcium neural activation experiments and PTH binding assay

CaCl_2_ (12 mM, Sigma, C5670), hPTH (1-34) (20 μg/mL, MCE, 52232-67-4), EDTA-Na2 (12 mM, Sigma, E9884), hPTH (1-34)-Lys (Biotin) (20 μg/mL, AnaSpec, AS-23647) and Biotin (1 μg/mL,Sigma, B4501) were injected intravenously into C57 mice. Animals were returned to their home cage and transcardially perfused for tissue collection after 90-120 mins.

### Immunofluorescence staining and *in situ* hybridization

Animals were anaesthetized with pentobarbital (100 mg/kg) and perfused intracardially with PBS followed with 4% PFA. Tissue was collected and fixed in 4% PFA at 4 °C for 48 hours, and then dehydrated with 30% sucrose solution for 72 hours. Coronal sections of brain (35 μm) and parathyroid glands (PTG, 15 μm) were cut using a Cryostat microtome (Leica, CM1950) at −20 °C. The sections were washed twice with PBS and blocked with blocking solutions (0.3% Tween20 in PBS with 5% goat normal serum or 5% bovine serum albumin) for 1 hour at room temperature, and then incubated overnight at 4 °C with appropriate primary antibodies diluted in 0.03% PBST. The primary antibodies used were: goat anti-Biotin (Sigma, F6762, 1:80), rabbit anticalcium sensing receptor (abeam, ab622l4, 1:500), mouse anti-CaMKII (abeam, ab22609, 1:500), goat anti-CD31 (R&D, AF3628, 1:20), rabbit anti-cFos (Cell signaling, #2250, 1:200), mouse anti-CGRP (abeam, 81887, 1:500), rabbit anti-ChAT (abeam, ab6168, 1:500), mouse anti cre-recombinase (Millipore, mab3120, 1:500), rabbit anti-GABA(Sigma, ab2052,1:500), chicken anti-GFP (abeam, abl3970,1:1000), rabbit anti-PTH (LSBio, LS-C191152, 1:500), rabbit anti-PTHIR (abeam, abl76393, 1:200), rabbit anti-PTH2R (abeam, abl 88760, 1:200), mouse anti-TH (Sigma, T1299, 1:500), and chicken anti-NeuN (Millipore, ABN9l, 1:500). Primary antibodies were washed three times with 0.1% PBST, and then replaced with appropriate secondary antibodies for 1 hour at room temperature and washed three times with 0.1% PBST. The secondary antibodies used were: Goat anti rabbit Alexa Fluor 488/594 (Jackson laboratory, 111-547-003/111-587-003, 1:200), Goat anti mouse Alexa Fluor 488/594 (Jackson laboratory, 115-547-003/115-587-003, 1:200), Anti-Goat IgG-FITC (Sigma, F7367, 1:400), Donkey anti rabbit Alexa Fluor 647 (abeam, ab150075, 1:1000), and Donkey anti goat Alexa Fluor 647 (abeam, abl 50131, 1:1000). Sections were then incubated with 0.5 μg/ml 4’,6-diamidino-2-phenylindole, dihydrochloride (DAPI; ThermoFisher, D1306) for 1 min and Fluoromount-G (Southern Biotech, 0100-01).

*In situ* hybridization was performed to localize GABAergic and glutamatergic neurons with *Vglut2, Gad1* and *Gad2* probes. The procedure of *in situ* hybridization followed the procedure described by Kondoh et al.^49^. Quantification of cell numbers labeled with different markers within different brain nuclei were calculated using 3 different slices from 3-4 different animals based on DAPI expression. The average of the three slices were taken and shown in graphs.

### Biochemical analysis

Mouse and rat serum PTH levels were assessed with mouse and rat PTH ELISA kits (LSBio, LS-5549/LS-5548). Total serum calcium levels were obtained using a colorimetric assay (Sigma, MAK022). The results were read by a Nano Quant plate reader (Tecan, Infinite 200Pro).

### Behavioral tests

Anxiety levels were assessed by an open-field test (OFT). In the OFT, mice were placed in a comer of a square field of 80 × 80 cm, with walls of 80 cm and sufficient illumination. The movement of animals from the entrance of the field/maze was recorded in an isolated room without interruption. The recording time for the OFT was 600 s. The time that each animal spent in different areas was assessed by analyzing software (Anymaze) to evaluate anxiety levels. Tail suspension tests (TST) were performed using a tail suspension cage (Bioseb, US). Mice were suspended by their tail with tape on the force transducer and the recording time was 360 s. The time spent immobile was analyzed using Anymaze within the Bioseb’s suspension cage.

Fluid consumption tests were performed in cages with two bottles. Animals were single housed to acclimatize to two bottles of water for 3 days and then completely water restricted for 12 hours before the test. Two bottles of water and 300 mM NaCl solution were provided to the animals during the test. Bottles were weighed every 2 hours within the first 6 hours during which the animals were free to access the fluids. The position of the bottles was changed after the first 1 hour to avoid position preference.

### Calcium tolerance test

Animal blood samples were collected before and after administration of CaCl_2_ *(i.p.* 3 μM for mice, 30 μM for rats) and serum PTH and calcium levels were assessed before, 5 min, 15 min and 30 min after administration. Blood was collected from the retro-orbital sinus of mice and from the jugular vein from rats under anesthesia with isoflurane.

### Micro-CT scanning and analysis

Mice femur and tibiae were collected and immersed in 4% paraformaldehyde before micro-CT scanning (SkyScan, model 1076). Scanning was performed using the following settings: isotropic voxel 11.53 μm, voltage 48 kV, current 179 μA, and exposure time of 1800 ms. Three-dimensional (3D) reconstruction was conducted using SkyScan NRecon software (version 1.6.8.0, SkyScan) with a voxel size of 8.66 μm. Datasets were reoriented using DataViewer (version 1.4.4.0, SkyScan), while the calculation of morphological parameters was carried out with the CTAn software (version 1.13.2.1, SkyScan). The 3D reconstructed models were displayed by CTVol software (version 2.2.3.0, SkyScan).

Trabecular bone was selected 0.1-0.9 mm distal to the proximal tibia growth plate. The region of interest (ROI) was selected from 2D images slice-by-slice by hand to exclude the cortical area and then binarization of the images with a global thresholding of gray level (70-255) as mineralized tissue, according to the tuning 3D reconstruction of the mineralized tissue. A Gaussian filter (radius=l) was used for 3D reconstruction. Quantitative analysis involved all bone areas within the ROI of the 3D images. Morphometric parameters included total volume (TV, m^3^), bone volume (BV, m^3^), bone fraction (BV/TV, m^3^), trabecular thickness (Tb. Th, 1/mm), trabecular number (Tb. N, 1/mm), and trabecular separation (Tb. SP, mm). In addition, the bone mineral density (BMD, g/cm^3^) of the whole trabecular bone was calibrated using the attenuation coefficient of two hydroxyapatite phantoms with defined mineral densities of 0.25 and 0.75 g/cm^3^.

Cortical bone was selected 6.5-7.2 mm proximal to the distal femur growth plate. The region of interest (ROI) was selected from 2D images with a threshold (85-255) as mineralized tissue. A Gaussian filter (radius=l) was used for 3D reconstruction. Quantitative analysis involved all bone areas within the ROI of the 3D images. Morphometric parameters included BV and BMD (within selected bone area).

### Electrophysiology

Animals were sacrificed under deep anesthesia and brains removed and placed in icecold cutting solution for 1 min (110 mM Choline Chloride, 2.5 mM KC1,0.5 mM CaCl_2_, 7 mM MgCl_2_, 1.3 mM NaH_2_PO_4_,1.3 mM Na-ascorbate, 0.6 mM Na-pyruvate, 20 mM glucose and 2.5 NaHCO_3_, 290-310 mOsm/kg, saturated with 95% O_2_ and 5% CO_2_). Coronal slices (300 μm) were cut using a vibrating microtome (VT1000, Leica), and then allowed to recover for 30 min at 37 °C in artificial cerebral spinal fluid (aCSF, 125 mM NaCl, 2.5 mM KC1,2 mM CaCl_2_,1.3 mM MgCl_2_, 1.3 mM NaH_2_PO_4_,1.3 mM Na-ascorbate, 0.6 mM Na-pyruvate, 5 mM glucose, 5mM sucrose and 2.5 mM NaHCO_3_, 300-310 mOsm/kg, saturated with 95% O_2_ and 5% CO_2_). After recovery, brain slices were transferred to a recording chamber and perfused with 2 ml/min aCSF. Patch pipettes were pulled from borosilicate glass (PG10150-4, World Precision Instruments) and filled with intrapipette solution (35 mM K-gluconate, 10 mM HEPES, 0.2 mM EGTA, 5 mM QX-314, 2 mM Mg-ATP, 0.1 mM Na-GTP, 8 mM NaCl, at 280~290 mOsm/kg and adjusted to pH 7.3 with KOH). Whole-cell patch clamp recording of SFO neurons was performed at room temperature (22-25 °C) with a Multiclamp 700B amplifier connected to a Digidata 1440A interface (Axon Instruments). Data were sampled at 10 kHz and analyzed with pClamp10 (Molecular Devices) or MATLAB (MathWorks).

### Ablation of the sympathetic nerves in the PTG

Sympathetic nervous ablation was performed with 6-hydroxydopamine hydrobromide (6-OHDA, Sigma, H116). Universal ablation of sympathetic nerves was performed in mice by *i.p.* injection of 15 mg/kg 6-OHDA. *In situ* ablation of sympathetic nerves in parathyroid glands of rats was performed by direct injection of 1 μL of 6-OHDA into the parathyroid glands. Calcium tolerance tests were performed 24 hours after 6-OHDA administration.

### Statistical analysis

In all studies, results were presented as mean ± standard error of the mean (SEM) with n= number of individual animals. Statistical analysis was performed using unpaired Student’s t-test for control and treatment comparisons, wherever appropriate, using the statistical program Prism version 7 (GraphPad Software, San Diego, CA, USA). A difference was accepted as statistically significant when probability (P) values were less than 0.05.

## Acknowledgements

This project was partly supported by the National Natural Science Foundation of China (82072489); Key Research Program of Frontier Sciences of Chinese Academy of Sciences (QYZDB-SSW-SMC056); Shenzhen Governmental Basic Research Grant (JC YJ20180507182301299).

## Author contributions

Fan Yang and William W Lu supervised the study. Lu Zhang, Nian Liu designed the experiments and conducted chemogenetic experiments, animal behavior studies, serum biochemical analysis and analyzed data. Jie Shao performed electrophysiological recordings in brain slices. Dashuang Gao performed PRV retrograde tracing experiments. Yunhui Liu performed sympathetic nerve ablation test.

---

**Supplementary figure 1.**
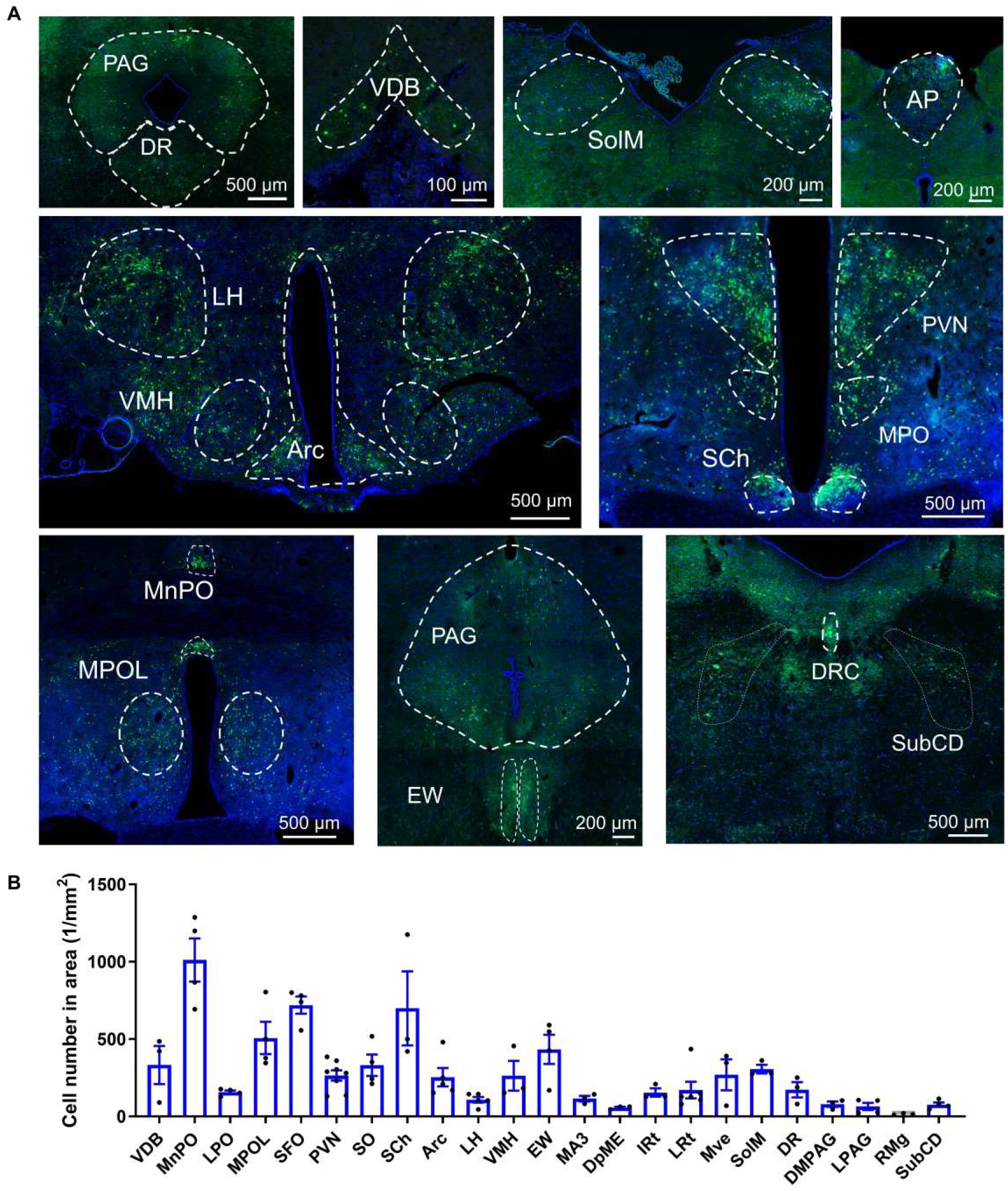
Poly-synaptic retrograde PRV labeling of specific rat brain nuclei. **A,** Pseudorabies virus (PRV) retrograde tracing from rat PTG to different brain nuclei, including: the AP, area postrema; Arc, arcuate nucleus; DMPAG, dorsomedial periaqueductal gray; DpME, deep mesencephalic nucleus; DR, dorsal raphe nucleus; EW, edinger-westphal nucleus; IRt, intermediate reticular nucleus; LH, lateral hypothalamic area; LPAG, lateral periaqueductal gray; LPO, lateral preoptic area; MA3, medial accessory oculomotor nucleus; MnPO, median preoptic nucleus; MPOL, medial preoptic nucleus, lateral part; Mve, medial vestibular nucleus; PVN, paraventricular thalamic nucleus; RMg, raphe magnus nucleus; SCh, suprachiasmatic nucleus, lateral part; SFO, subfornical organ; SO, supraoptic nucleus; SolM, nucleus of the solitary tract, medial part; SubCD, subcoeruleus nucleus, dorsal part; VDB, nucleus of the vertical limb of the diagonal band; VMH, ventromedial hypothalamic nucleus. N=3 rats. **B,** Statistical analysis of signal density of different brain nuclei 7 days after PRV injection from rat PTG. All error bars and shaded areas show mean ± s.e.m.

**Supplementary figure 2.**
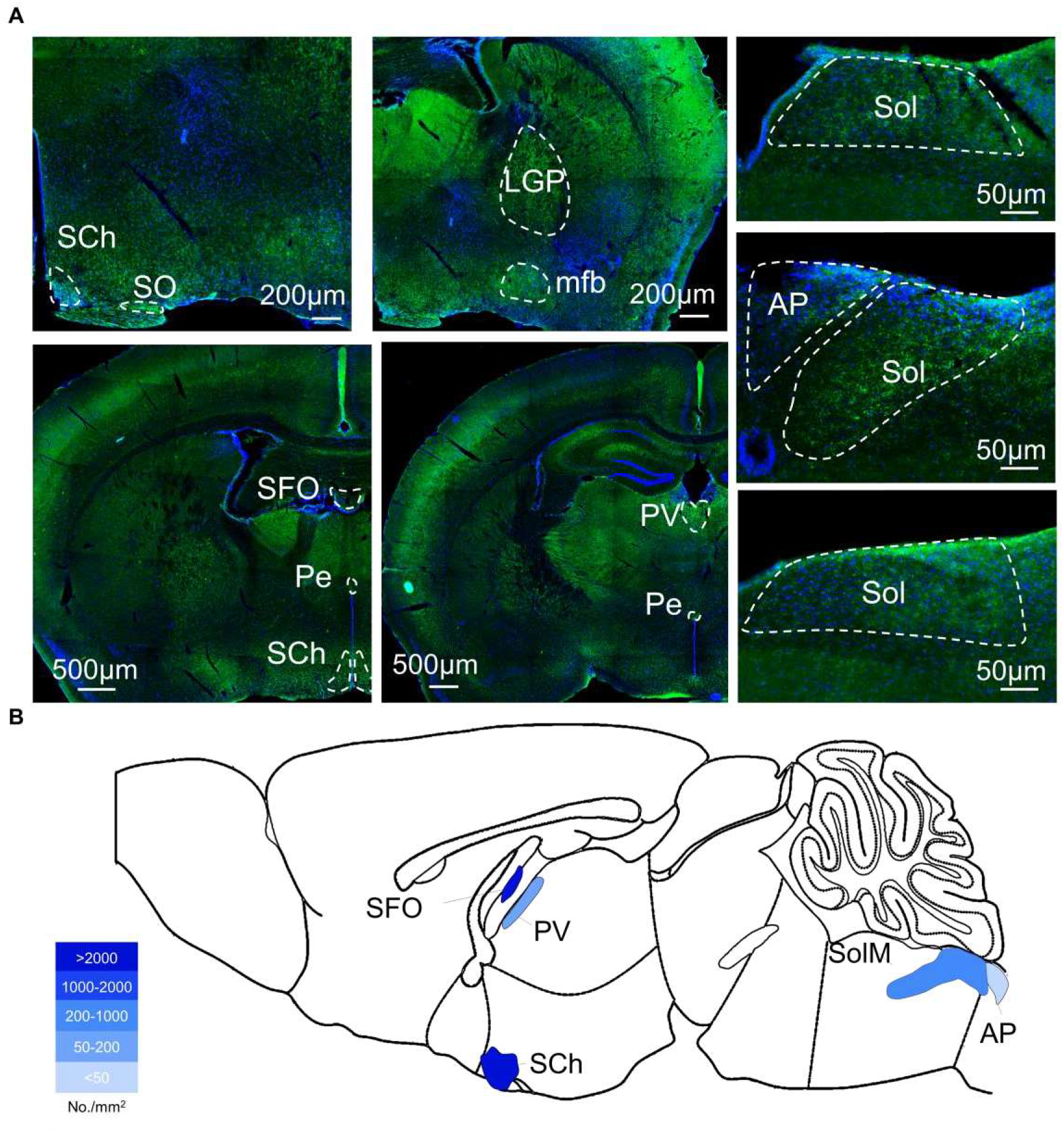
PTH-biotin binding to specific brain nuclei after peripheral *i.v.* administration. **A,** Immunofluorescence staining of biotin indicating the PTH-biotin binds to different nuclei in the C57 mouse brain, including: the AP, area postrema; LGP, lateral globus pallidus; mfb, medial forebrain bundle; Pe, periventricular hypothalamic nucleus; PV, periventricular fiber system; SCh, suprachiasmatic nucleus; SFO, subfornical organ; SO, supraoptic nucleus; Sol, solitary tract. **B**, PTH-biotin binding nuclei location and the signal density (n=3 mice).

**Supplementary figure 3.**
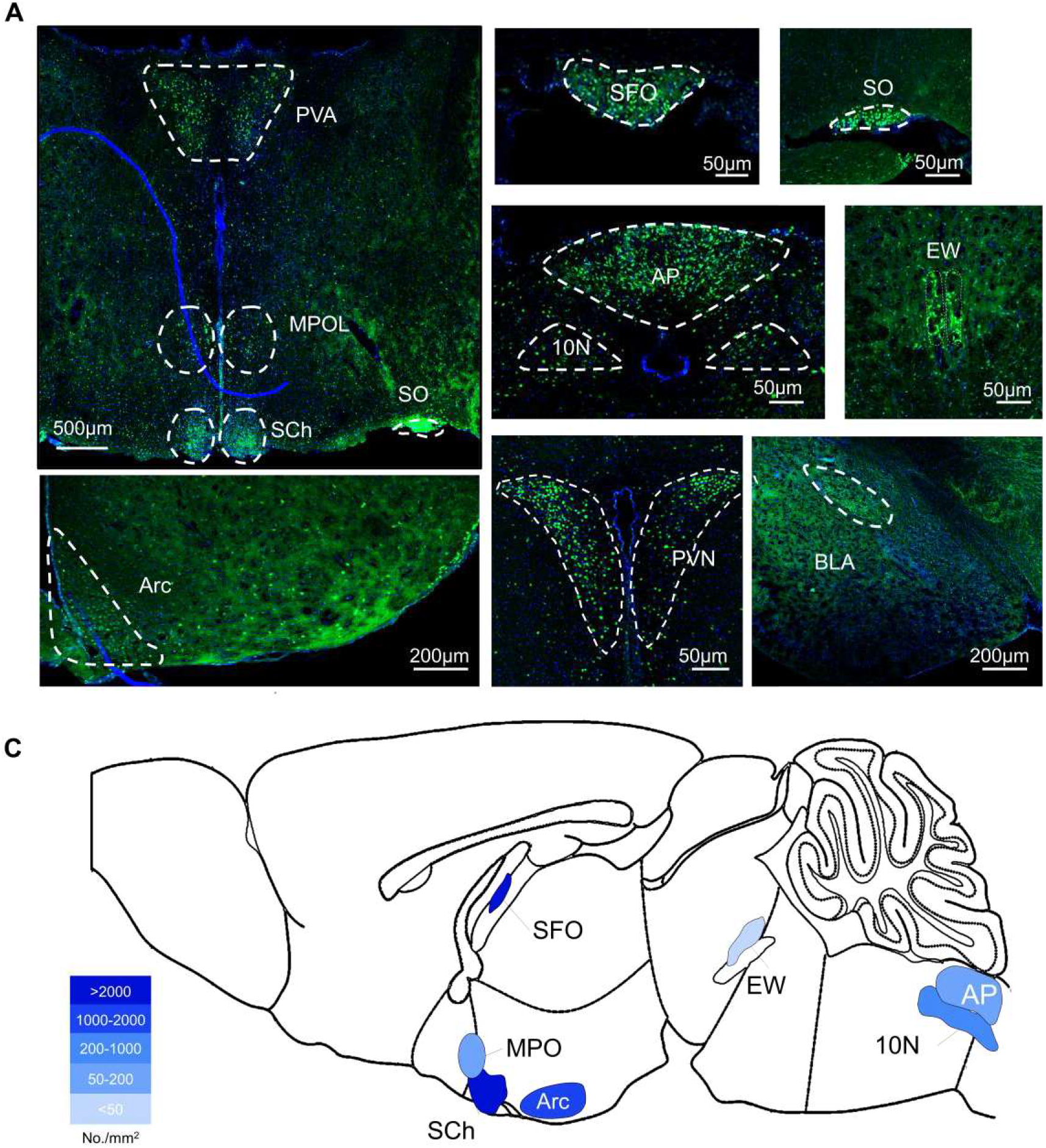
cFos expression in the mouse brain after peripheral ¿v. PTH stimulation. **A,** Immunofluorescence staining of cFos (green) were observed in specific brain nuclei including: the ION, dorsal motor nucleus of vagus; Arc, arcuate hypothalamic nucleus; AP, area postrema; BLA, basolateral amygdaloid nucleus, anteriorpart; EW, Edinger-Westphal nucleus; MPOL, medial preoptic nucleus, lateral part; PVA, paraventricular thalamic nucleus, anterior part; PVN, paraventricular thalamic nucleus; SCh, suprachiasmatic nucleus; SFO, subfornical organ; SO, supraoptic nucleus. **B**, Locations of PTH-responding nuclei and relative cFos expression density (n=3 mice).

**Supplementary figure 4.**
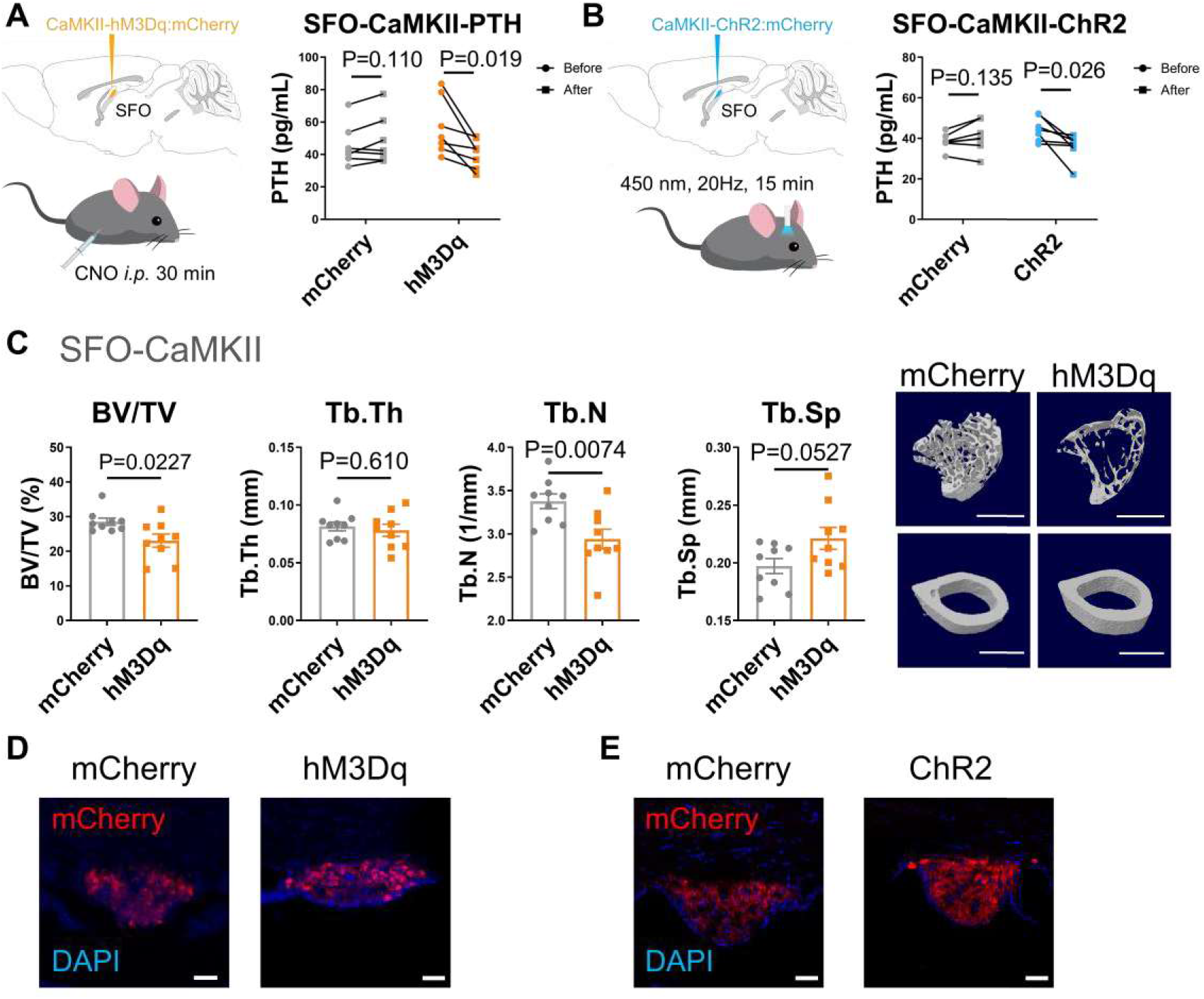
Rapid serum PTH changes, bone phenotype and behavior analysis following stimulation of SFO^CaMKII^ neurons. **A,** Serum PTH changes after chemogenetic stimulation of CaMKII neurons (before and 30 min after CNO *i.p.* injection) in SFO of C57 mice (n=7). B, Serum PTH changes after optogenetic stimulation of CaMKII neurons in SFO of C57 mice (n=6-7). **C,** Bone analysis after stimulation of SFO^CaMKII^ neurons. ***Left bar charts,*** tibia trabecular percent bone volume (BV/TV), trabecular thickness (Tb.Th), trabecular number (Tb.N), trabecular separation (Tb.Sp) assessed by MicroCT (n=9 mice). ***Right,*** 3D reconstruction of MicroCT analysis of tibia trabecular and femur cortical bones in hM3Dq and mCherry groups. Scale bar, 1 mm. **D,** The expression of hM3Dq-mCherry in CaMKII neurons in SFO. Scale bar, 50 μm. E, The expression of ChR2-mCherry in CaMKII neurons in SFO. Scale bar, 50 μm. P analyzed by paired two-tailed t-test in **A-B;** P analyzed by unpaired two-tailed t-test in **C.** All error bars and shaded areas show mean ± s.e.m.

**Supplementary figure 5.**
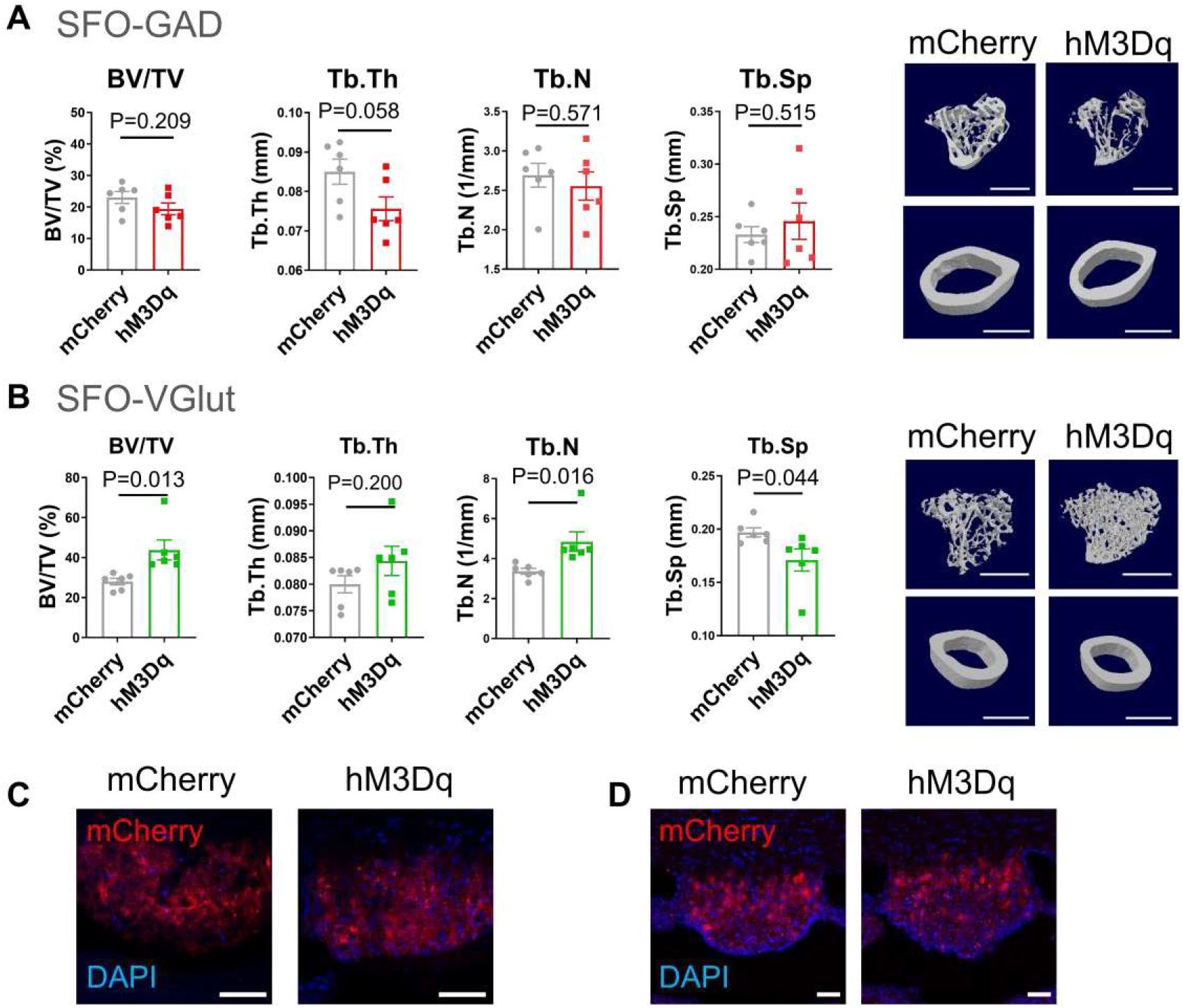
Bone analysis following chemogenetic regulation of SFO^GAD^ and SFO^VGlut^ neurons. **A**, Bone analysis after chronic chemogenetic stimulation of SFO^GAD^ neurons for 4 weeks. ***Left bar charts,*** tibia trabecular percent bone volume (BV/TV), trabecular thickness (Tb.Th), trabecular number (Tb.N), trabecular separation (Tb.Sp) assessed by MicroCT analysis (n=6 mice). ***Right,*** 3D reconstruction of MicroCT analysis of tibia trabecular and femur cortical bones in hM3Dq and mCherry groups. Scale bar, 1 mm. **B**, Bone analysis after chemogenetic stimulating SFO^VGlut^ neurons for 4 weeks. ***Left bar charts,*** tibia trabecular percent bone volume (BV/TV), trabecular thickness (Tb.Th), trabecular number (Tb.N), trabecular separation (Tb.Sp) assessed by MicroCT analysis (n=6 mice). ***Right,*** 3D reconstruction of MicroCT analysis of tibia trabecular and femur cortical bones in hM3Dq and mCherry groups of VGlut2-cre mice. Scale bar, 1 mm. **C,** The expression of hM3Dq-mCherry in GABAergic neurons of SFO. Scale bar, 50 μm. **D,** The expression of hM3Dq-mCherry in glutamatergic neurons in SFO. of SFO. Scale bar, 50 μm. P analyzed by unpaired two-tailed t-test. All error bars and shaded areas show mean ± s.e.m.

**Supplementary figure 6.**
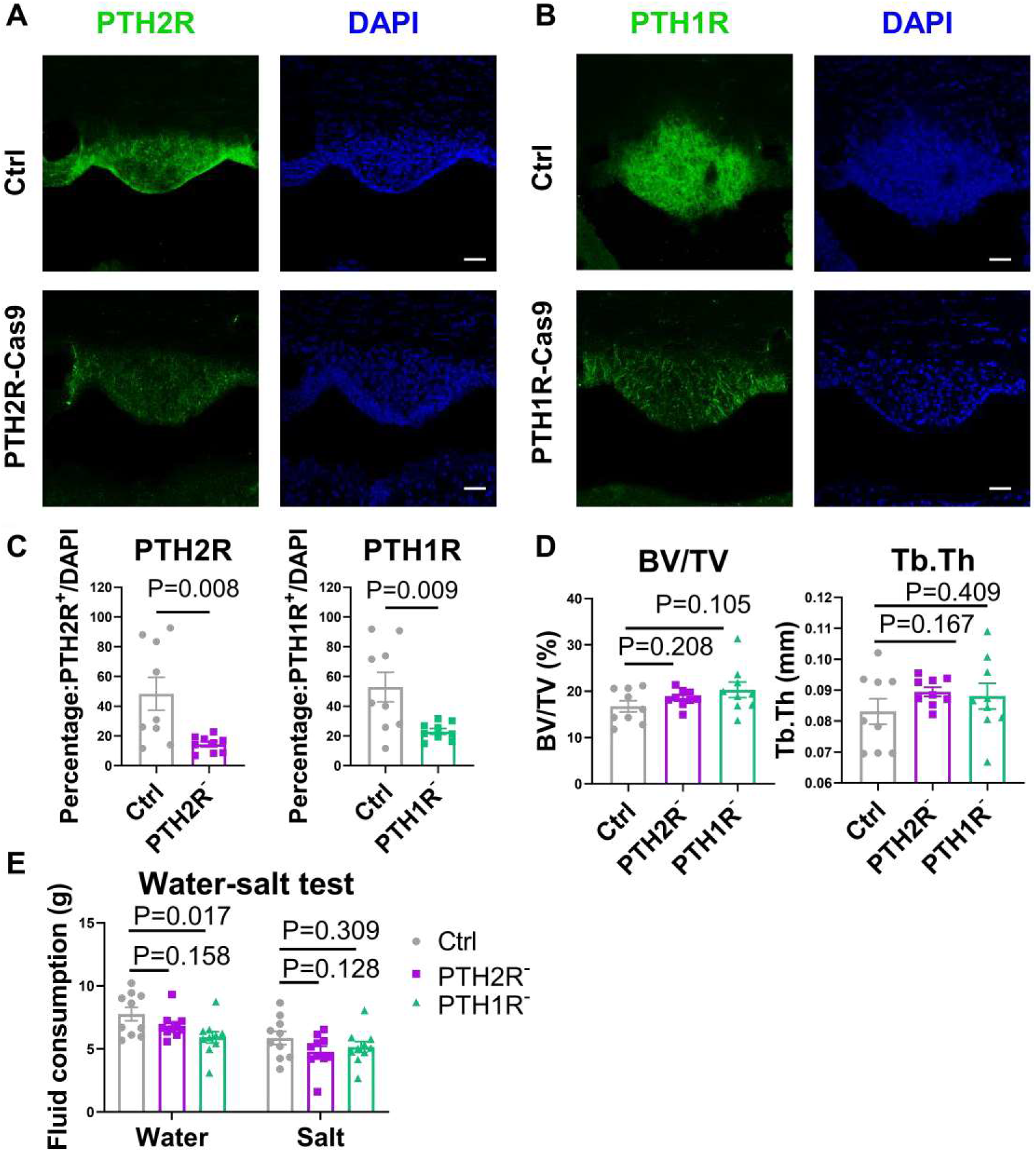
Knockdown of PTH1R/PTH2R expression in SFO and related bone changes. **A,** Immunofluorescence staining of PTH2R in SFO of control and PTH2R knockdown mice. Scale bar, 50 μm. **B**, Immunofluorescence staining of PTH1R in SFO of control and PTH1R knockdown mice. Scale bar, 50 μm. **C**, Quantification of PTH2R/PTH1R positive cells in control and knockdown group (n=9 mice). **D**, Percentage of Bone volume (BV/TV), trabecular thickness (Tb.Th) of C57 mice tibia after knockdown of PTH1R/PTH2R (n=9 mice). **E**, Fluid consumption (water or 0.3M NaCl) tests in C57 mice after knockdown of PTH1R/PTH2R expression in the SFO (n=9 mice). P analyzed using unpaired two-tailed t-tests. All error bars and shaded areas show mean ± s.e.m.

**Supplementary figure 7.**
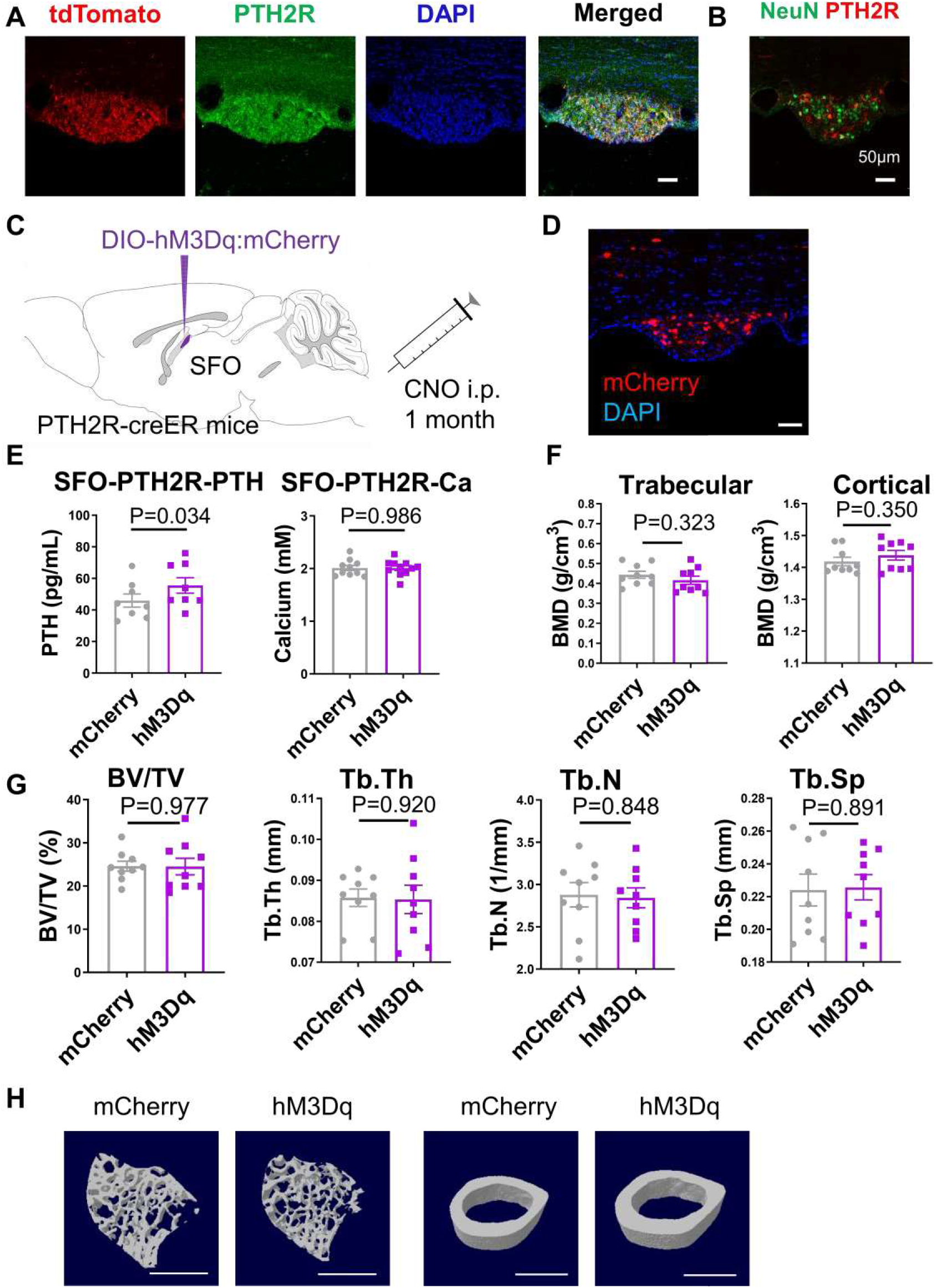
Serum PTH and bone metabolism changes after chemogenetic stimulation of SFO^PTH2R^ neurons in PTH2R-creER^12^ mice. **A,** Immunofluorescence (IF) staining of PTH2R colocalized with tdTomato signal in SFO of PTH2R-CreER^T2^ X Ail4 mice. Scale bar, 50 μm. **B**, PTH2R neurons labeled by AAV-DIO-mCherry transduction to the SFO of PTH2R-CreER^17^ mice and colocalization with IF staining of NeuN (green). Scale bar, 50 μm. **(C-G),** Chemogenetic activation of PTH2R neurons in the SFO of PTH2R-CreER^T2^ mice (n=8-9 mice). **C,** Schematic showing injection of AAV-DIO-hM3Dq/mCherry into the SFO of PTH2R-CreER^T2^ mice followed with 1-month CNO *i.p.* administration. **D,** Expression of hM3Dq-mCherry within PTH2R neurons in SFO. Scale bar, 50 μm. **E,** Serum PTH, total calcium level of PTH2R-creER^T2^ mice after chemogenetic stimulation of SFO^PTH2R^ neurons. **F,** Tibia trabecular bone mineral density (BMD), and femur cortical BMD from MicroCT analysis of PTH2R-creER^T2^ mice after chemogenetic stimulation of SFO^PTH2R^ neurons. **G**, Percentage of bone volume (BV/TV), trabecular thickness (Tb.Th), trabecular numbers (Tb.N) and trabecular seperation (Tb.Sp) of PTH2R-creER^T2^ mice after chemogenetic stimulation of SFO^PTH2R^ neurons. **H**, 3D reconstruction of MicroCT analysis of tibia trabecular and femur cortical bones in mCherry and Hm3Dq groups. Scale bar, 1 mm. P analyzed using unpaired two-tailed t-test. All error bars and shaded areas show mean ± s.e.m.

**Supplementary figure 8.**
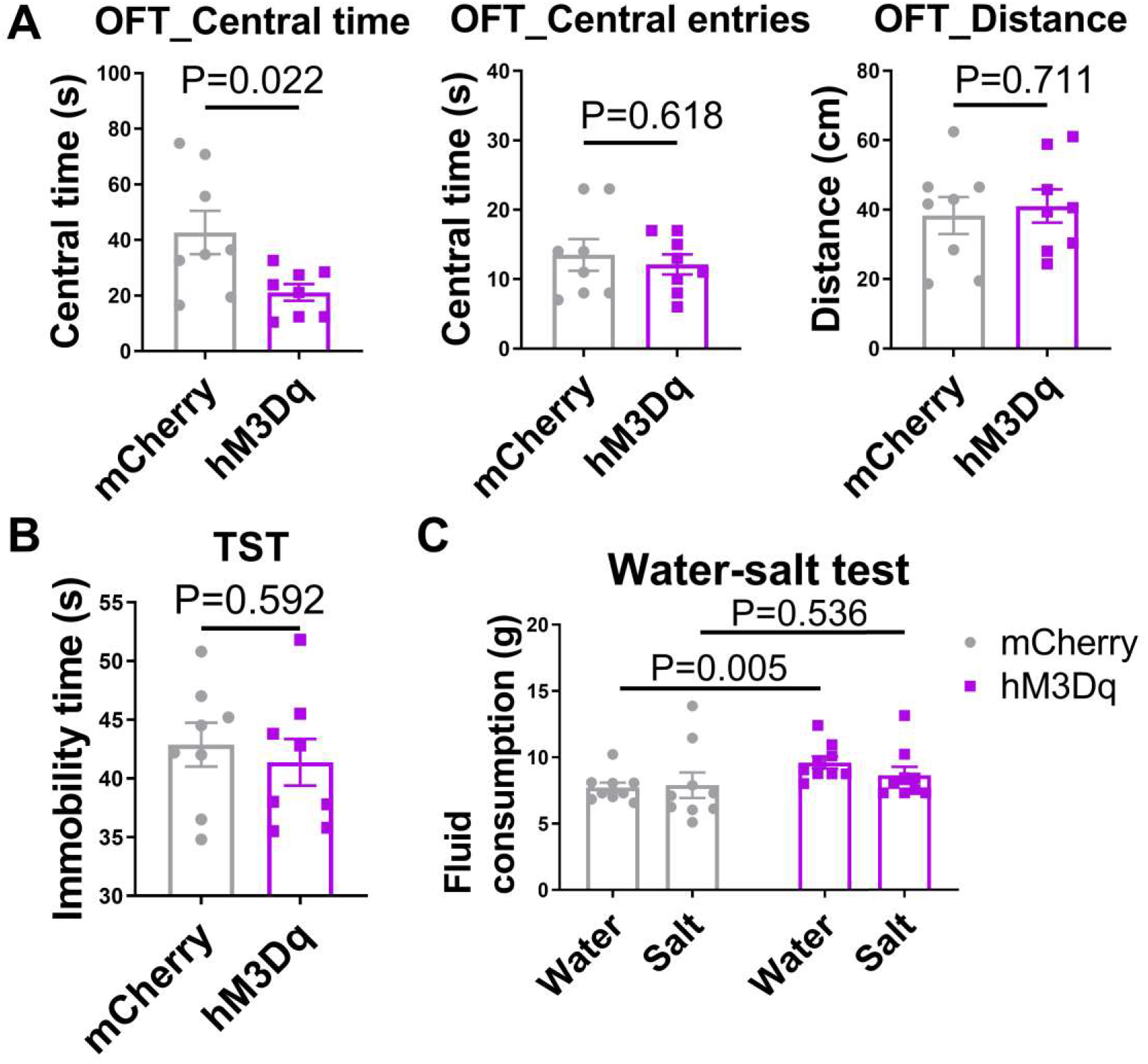
Behavior changes after chemogenetic stimulation of SFO^pth2r^ neurons in PTH2R-creER^T2^ mice. **A**, Time spent in center, central entries and total distance during open field test of PTH2R-creER^T2^ mice after chemogenetic stimulation of SFO^PTH2R^ neurons, (n=8 mice) **B**, Total immobility time during tail suspension test, (n=8 mice) **C,** Water and salt consumption of PTH2R-creER^T2^ mice after chemogenetic stimulation of SFO^PTH2R^ neurons, (n=8 mice) P analyzed using unpaired two-tailed t-test. All error bars and shaded areas show mean ± s.e.m.

**Supplementary figure 9.**
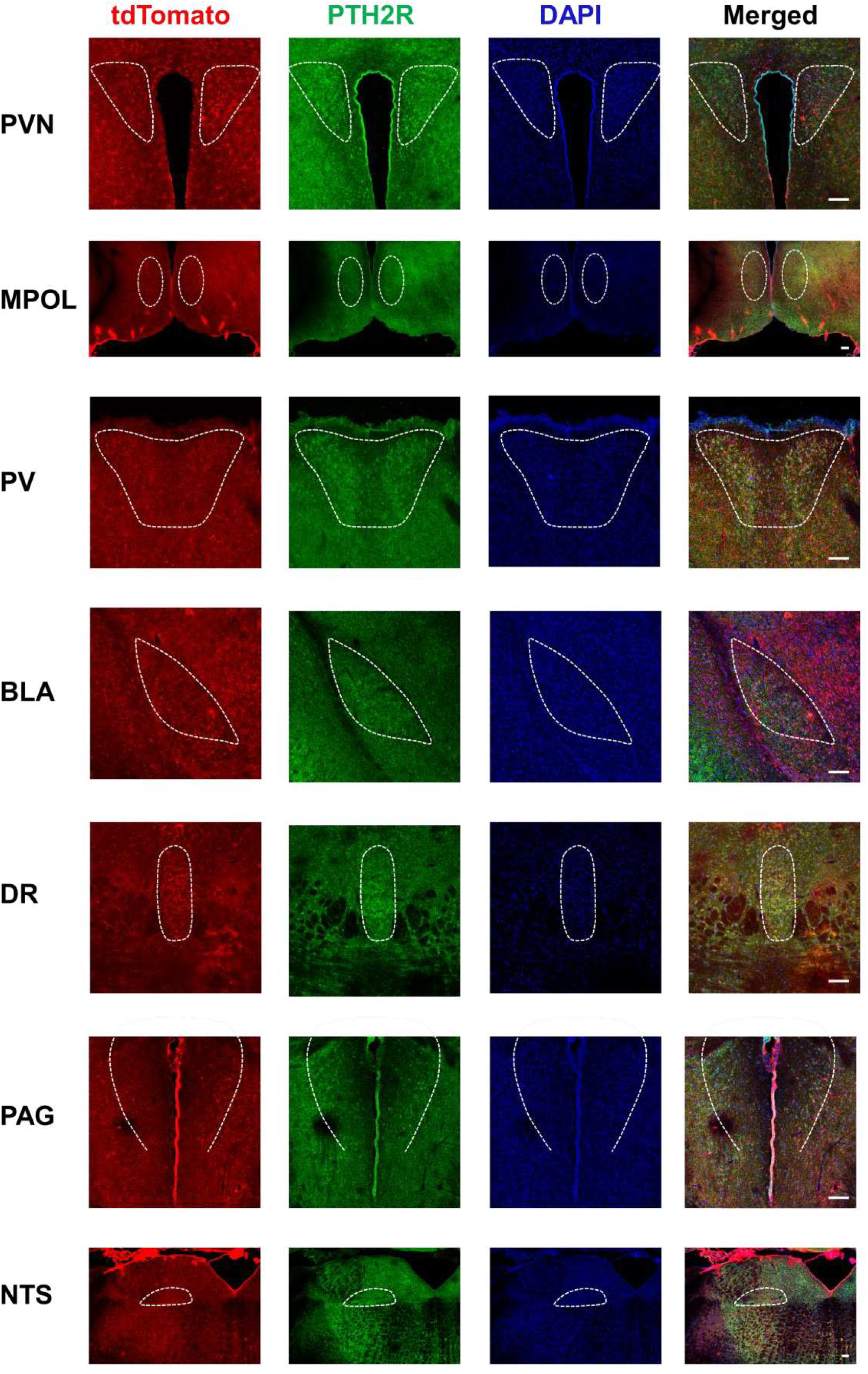
Colocalization of immunofluorescence staining of PTH2R with tdTomato signals in PTH2R-CreER^T2^ X Ai14 mice brain. Colocalization of immunofluorescence staining of PTH2R and tdTomato signals in PTH2R-creER^τ2^ X Ai14 mice. PVN, paraventricular nucleus; MPOL, medial preoptic nucleus, lateral part; PV, periventricular nucleus; BL A, basolateral amygdala; DR, dorsal raphe; PAG, periaqueductal gray; NTS, nucleus of the solitary tract; Arc, arcuate nucleus. Scale bar, 100μm.

**Supplementary figure 10.**
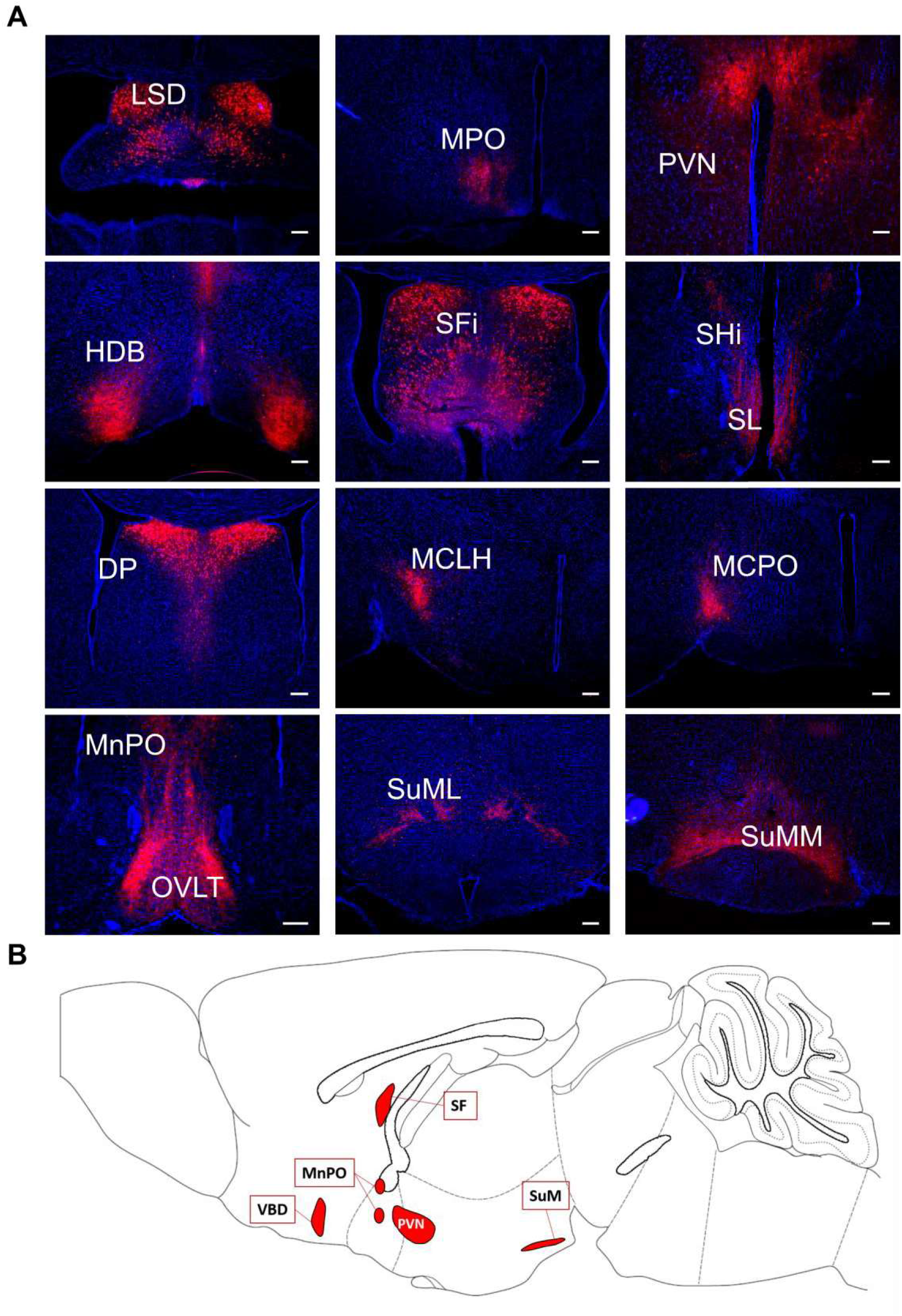
Neural projections of SFO^CaMKII^ neurons into different brain nuclei in C57 mice. **A,** SFO^CaMKII^ neurons labeled with AAV-CaMKII-mCherry project terminals into brain nuclei, including: the DP, dorsal peduncular cortex; HDB, nucleus of the horizontal limb of the diagonal band; LSD, lateral septal nucleus, dorsal part; MCLH, magnocellular nucleus of the lateral hypothalamus; MCPO, magnocellular preoptic nucleus; MnPO, median preoptic nucleus; MPO, medial preoptic nucleus; OVLT, vascular organ of lamina terminalis; PVN, paraventricular thalamic nucleus; SFi, septofimbrial nucleus; SHi, septohippocampal nucleus; SL, semilunar nucleus; SuML, supramammillary nucleus, lateral part; SuMM, supramammillary nucleus, medial part. Scale bar, 50μm B, Locations of brain nuclei receiving high density SFO^CaMKII^ neuronal projections (n=3 mice).

**Supplementary figure 11.**
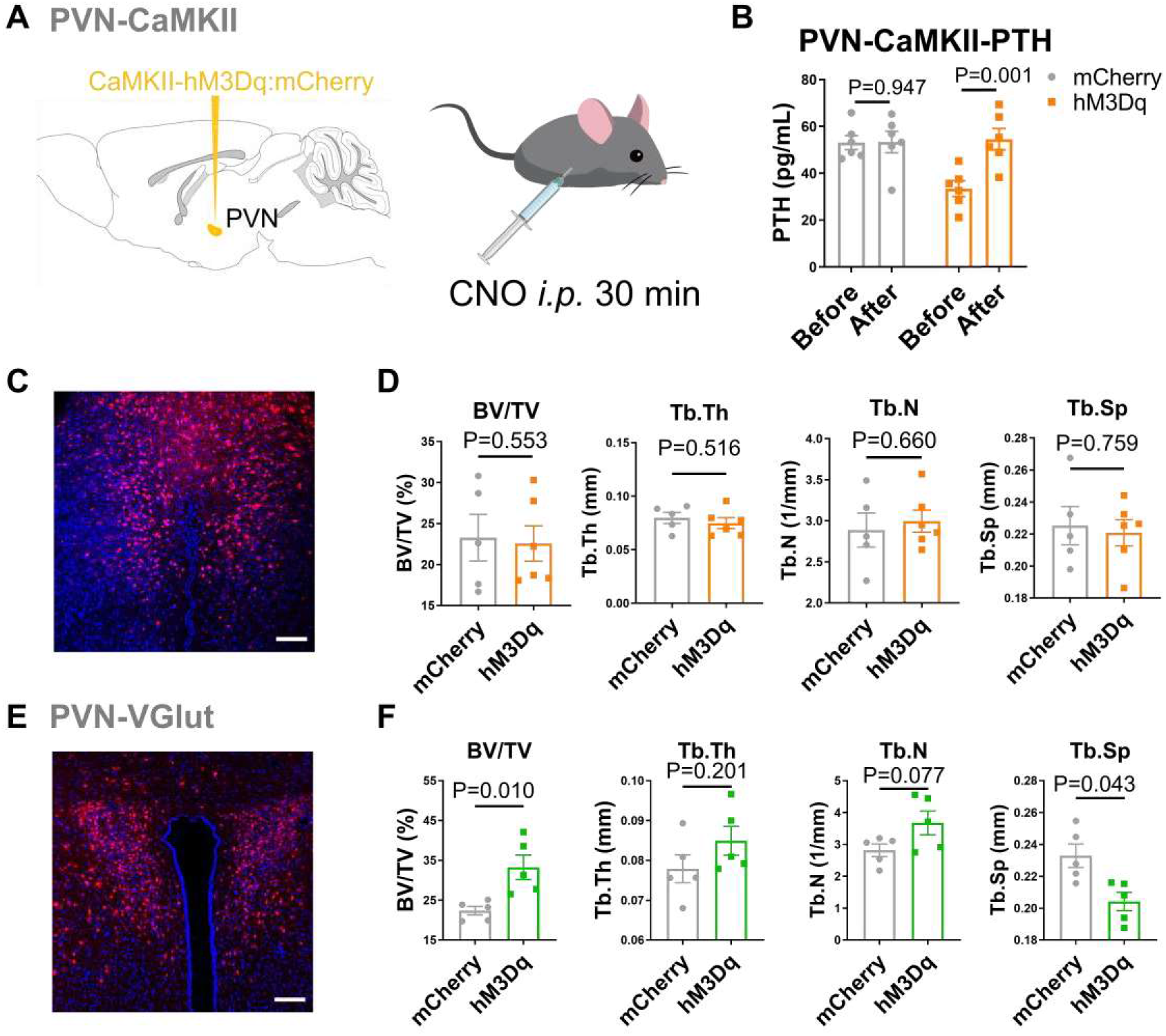
Serum PTH, bone structure and behavior changes after chemogenetic stimulation of PVN^CaMKΠ^ and PVN^VGlut^ neurons. **A,** Schematic showing the injection of AAV-CaMKII-hM3Dq/mCherry in to PVN OF C57 mice and CNO administration. Serum are collected 30 min after CNO administration. **B**, Rapid Serum PTH changes after chemogenetic stimulation of CaMKII neurons (before and 30 min after CNO *i.p.* injection) in the PVN of C57 mice (n=5). **C,** Expression of hM3Dq-mCherry in CaMKII neurons in PVN. Scale bar, 100 μm. **D**, The tibia trabecular percent bone volume (BV/TV), trabecular thickness (Tb.Th), trabecular numbers (Tb.N), trabecular separation (Tb.Sp) assessed by MicroCT analysis after chemogenetic stimulation of PVN^CaMK[I^ neurons for 1 month in C57 mice (n=5-6 mice). **E,** Expression of hM3Dq-mCherry in glutamatergic neurons in PVN. Scale bar, 100 μm. **F**, The tibia trabecular percent bone volume (BV/TV), trabecular thickness (Tb.Th), trabecular numbers (Tb.N), trabecular separation (Tb.Sp) assessed by MicroCT analysis after chemogenetic stimulation of PVN^VGIul^ neurons for 1 month in VGlut2-cre mice (n=5 mice). P analyzed using unpaired two-tailed t-test. All error bars and shaded areas show mean ± s.e.m.

**Supplementary figure 12.**
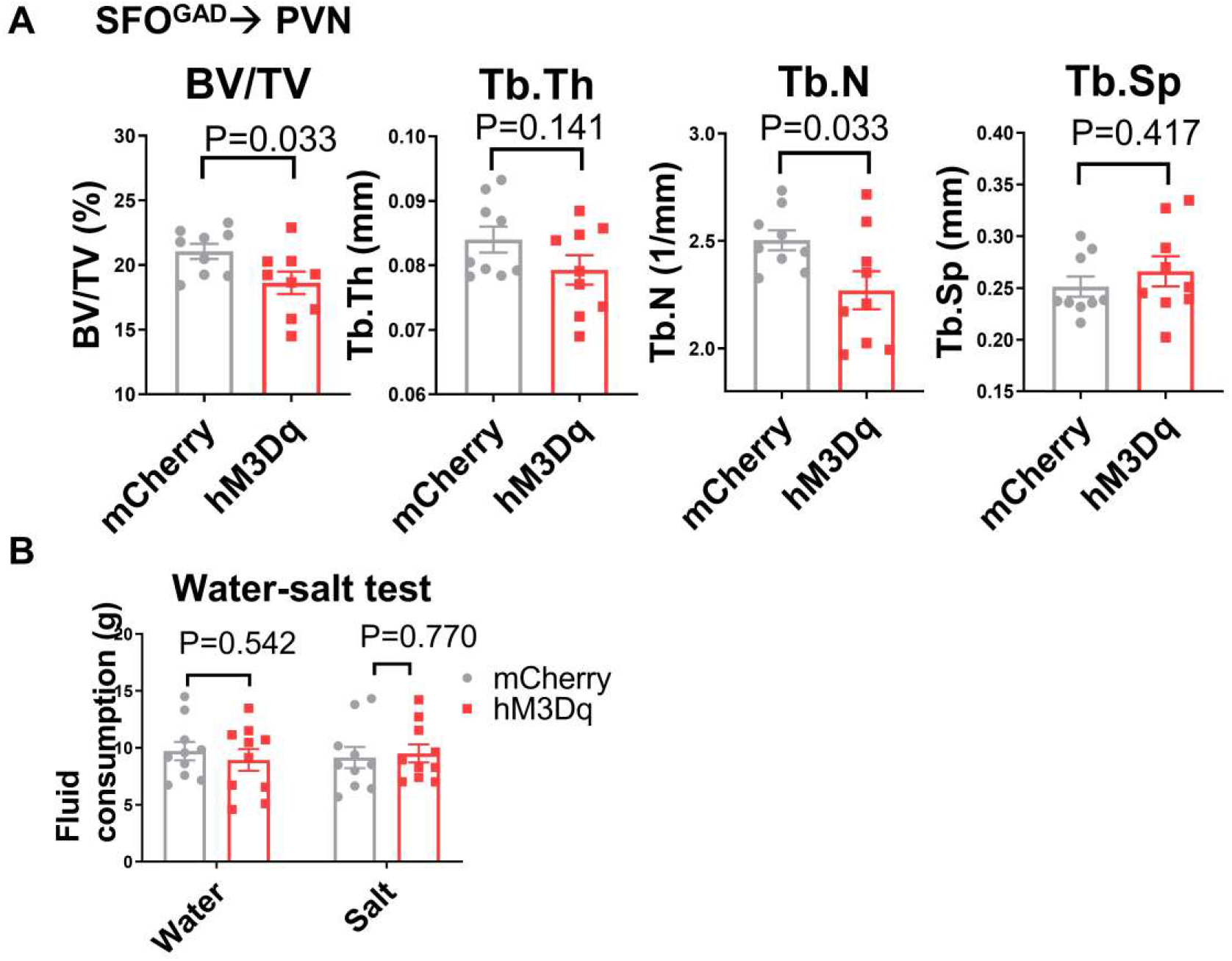
Bone analysis and behavioral tests following stimulation of the SFO^GAD^→PVN neural circuit. **A,** Tibia trabecular percent bone volume (BV/TV), trabecular thickness (Tb.Th), trabecular numbers (Tb.N), trabecular separation (Tb.Sp) assessed by MicroCT analysis after chemogenetic stimulation of SFO^GAD^→PVN for 1 month in C57 mice (n=9 mice). **B**, Water and salt consumption of C57 mice after chemogenetic stimulation SFO^GAD^→PVN for 1 month (n=lO mice). P analyzed using unpaired two-tailed t-test. All error bars and shaded areas show mean ± s.e.m.

**Supplementary figure 13.**
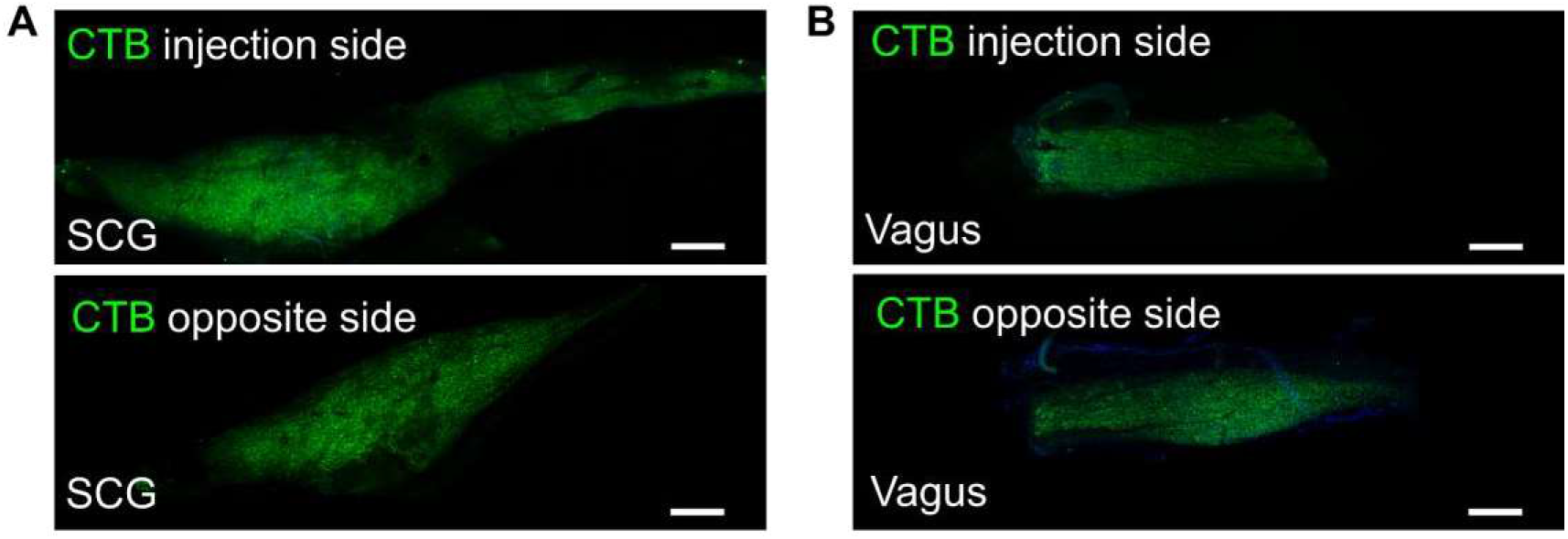
Neural specific retrograde tracing from rat parathyroid glands to upstream ganglia. **A,** Choleral toxin B (CTB) signals observed in both the injection side and opposite side of sympathetic superior cervical ganglia (SCG). Scale bar, 500 μm. **B**, CTB signals observed in both the injection side and opposite side of sympathetic inferior vagus ganglia (Vagus). Scale bar, 500 μm.

**Supplementary figure 14.**
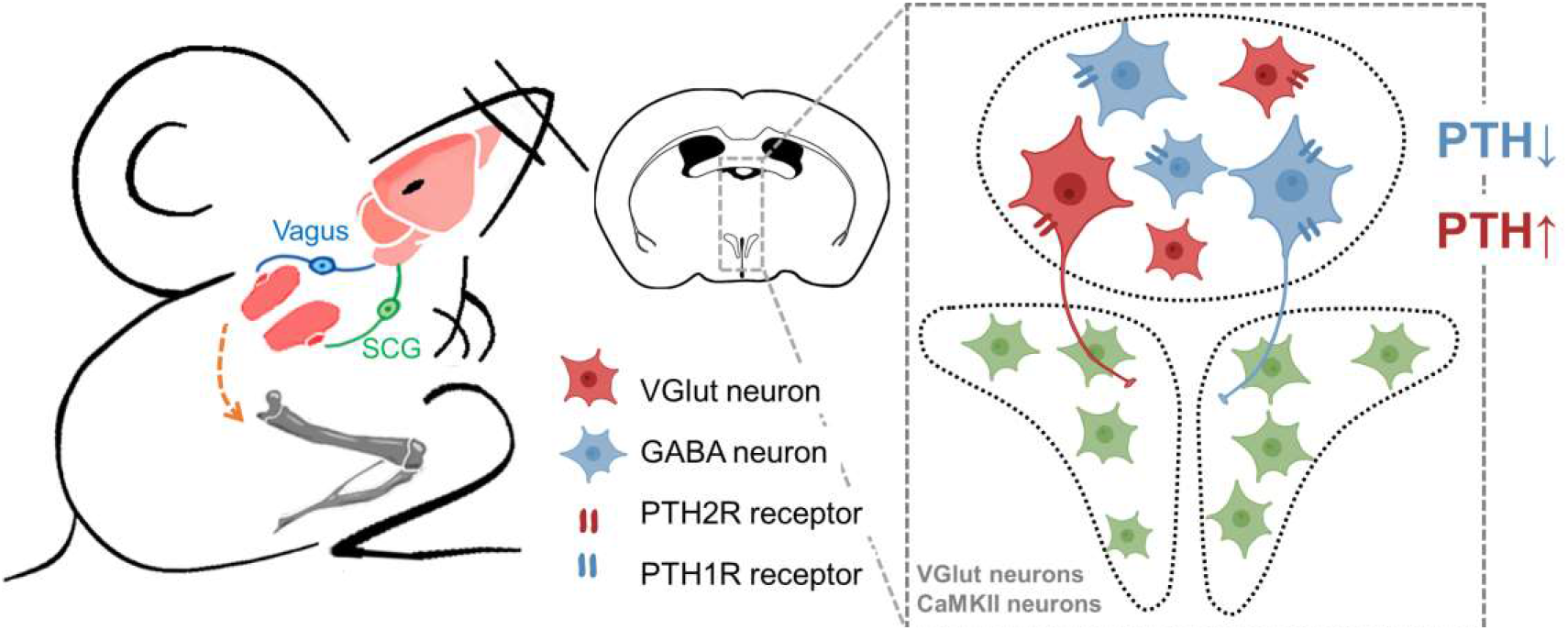
Schematic summarizing the mechanism underlying the neural 2 regulation of PTH levels in the SFO and PVN. PTH, parathyroid hormone; PTH1R, parathyroid hormone receptor 1; PTH2R, parathyroid hormone receptor 2; SCG, superior cervical ganglia; VGlut, vesicular glutamate transporter; GABA, gamma-Aminobutyric acid.

